# singIST: an integrative method for comparative single-cell transcriptomics between disease models and humans

**DOI:** 10.1101/2024.12.20.629624

**Authors:** Aitor Moruno-Cuenca, Sergio Picart-Armada, Alexandre Perera-Lluna, Francesc Fernández-Albert

## Abstract

**Motivation:** Disease models serve as fundamental tools in drug discovery and early-stage drug development. However, these models are not a perfect reflection of human disease, and selecting a suitable model can be challenging. Existing computational approaches for molecular validation of pathophysiological resemblance to human conditions at single-cell resolution remain limited. Although quantitative computational methods exist to inform this selection, they are very limited at the single-cell resolution, which can be critical for model selection. Quantifying the resemblance of disease models to the human condition with single-cell technologies in an explainable, integrative, and generalizable manner remains a significant challenge.

**Results:** We present singIST, a computational method for comparative single-cell transcriptomics analysis between disease models and human conditions. singIST provides explainable quantitative measures on disease model similarity to human condition at both pathway and cell type levels, highlighting the importance of each gene in the latter. These measures account for orthology, cell type presence in the disease model, cell type and gene importance in human condition, and gene changes in the disease model measured as fold change. This is achieved within a unifying framework that controls for the intrinsic complexities of single-cell data. We tested our method using three well-characterized murine models of moderate-to-severe Atopic Dermatitis, demonstrating its ability to recapitulate established biological knowledge while generating novel hypothesis through pathway-level analysis.

**Availability and implementation:** R library at https://github.com/DataScienceRD-Almirall/singIST and implementation code at https://github.com/DataScienceRD-Almirall/singIST_paper_results

**Author Summary:** In this study, we set out a method to improve how researchers evaluate disease models - tools that are essential for understanding human illness and developing new treatments. A major limitation of current methods is that they overlook the fine-scale differences between disease models and human conditions at the single-cell level. Yet, many diseases are driven by changes that occur in specific cell types.

To address this, we developed singIST, a method that allows us to compare disease models to human diseases using single-cell gene expression data. What makes singIST unique is its ability to provide clear, interpretable measures of similarity between a disease model and the human condition, specifically at the level of cell types and groups of functionally related genes.

We demonstrated the method using three well-characterized mouse models of atopic dermatitis and showed that singIST could both reflect known disease biology and uncover new insights into disease mechanisms. Further, we developed statistical tests to validate the trained models. Our approach is flexible and broadly applicable, making it a valuable tool for improving the selection and understanding of disease models in biomedical research.

## 1. Introduction

Disease models are biological experimental systems to study human disease. These models are designed to mimic the pathophysiology, progression, and response to treatments observed in human conditions. These models serve as the backbone to drug discovery and early drug development activities; drug target validation and characterization (Emmerich et al., 2021); compound screening (Elitt et al., 2018; Wei et al., 2021); preclinical studies to identify a lead candidate from several targets, select optimal formulation, posology and route of administration (Shegokar, 2020); guide early phase clinical trial design (Steinmetz and Spack, 2009; Loewa et al., 2023). However, the validation of molecular physiology, etiology and pathogenesis of disease models to that of human condition remain a challenge, which contribute to high rates of drug development attrition (Storey et al., 2022).

Recently, there have been methodological advancements in bioinformatics to quantitatively assess the validity of disease models in mimicking a human condition, through bulk transcriptomics. Found In Translation (FIT) (Normand et al., 2018) is a statistical methodology, relying on regularized linear regression models, that leverages bulk transcriptomics data to extrapolate murine disease models’ gene expression to expression changes that would be equivalent in the human condition, by using disease models’ Fold Changes (FC). Another approach is In Silico Treatment (IST) (Picart-Armada et al., 2024), a computational method that assesses translation of disease-related bulk gene expression patterns between animal models and humans, by also simulating observed disease models’ FC onto humans, providing an interpretable measure of their transcriptomics similarity. Nonetheless, evaluating disease models using bulk transcriptomics methods may lack the necessary granularity to underpin changes in specific cell populations involved in the pathological manifestations of the human condition. This is particularly true for Immune-mediated inflammatory diseases (IMIDs), whose pathogenesis is primarily driven by lymphoid cells (McInnes and Gravallese, 2021; Pisetsky, 2023). FIT nor IST provide a trivial approach to accommodate for single-cell data. Current methodologies for comparative analysis of single-cell transcriptomic changes in disease models’ to that of human condition are scarce. A recurrent approach is to perform an Overlapping Differentially Expressed Genes (ODEGs) analysis between disease models and human condition (Kim et al., 2019; Li et al., 2023; Ali et al., 2024), yet ODEGs have been proven to be suboptimal as it treats every gene direction and magnitude as equal posing the need for more sophisticated approaches (Lawhorn et al., 2018). Another strategy is performing a dimensionality reduction technique (CCA, NNMF, tSNE) on disease models’ and human scRNA-seq data and compare the obtained latent factors (Gao et al., 2021; Karmele et al., 2023; Franzén et al., 2024), which poses difficulty in interpreting and quantifying the similarity between both.

To address the challenges in single-cell transcriptomics analysis, we introduce singIST, a flexible computational method built on the foundation of IST. singIST facilitates comparative analysis between disease models and human conditions by accounting for orthology, cell type agreement, adaptive sparsity, and the importance of genes and cell types. It provides interpretable measures of transcriptomic similarity at different levels of granularity. We demonstrate its potential by assessing well-characterized murine models of Atopic Dermatitis (AD), a skin IMID, focusing on deregulated biological pathways.

## 2. Materials and methods

### 2.1 Materials

#### 2.1.1 Human data

Human moderate to severe AD scRNA-seq data correspond to patients diagnosed with chronic AD in early childhood, obtained from all Healthy Control (HC) and AD skin suction blisters samples analyzed in Bangert et al. (2021). In total, 4 HC and 5 AD skin suction blisters were used. Metadata and GEO identifiers can be found in Supplementary Material S4. Cell type populations modelled are those identified by Bangert et al. (2021): T-cells, Melanocytes, Dendritic Cells, Langerhans Cells and Keratinocytes. Raw counts were pseudobulked and posteriorely log normalized using Seurat v5.0.1. Human scRNA-seq were obtained by 10x Genomics sequencing platform.

#### 2.1.2 Disease model data

We evaluate scRNA-seq of three epicutaneous sensitized murine models that cause and AD like eczema phenotype; Oxazolone (OXA) and Imiquimod 5% cream (IMQ), obtained from Liu et al. (2020); Ovalbumine (OVA), obtained from Leyva-Castillo et al. (2022), the latter two are also stablished murine models in Psoriasis and Asthma, respectively. Metadata and GEO identifiers can be found in Supplementary Material S4. All disease models, and their respective controls, have 3 replicates of ear skin biopsies. To allow for comparison with human data, the existing cell type annotations of disease models were mapped to match those observed in the human dataset, the mapping of this relation is shown in Supplementary Material S2. Likewise to human samples, disease model samples were pseudobulked and log normalized thereafter. All disease models scRNA-seq were obtained by 10x Genomics sequencing platform.

#### 2.1.3 Pathway data

Pathways under analysis were selected from Brunner et al. (2017), all enriched pathways in human moderate to severe AD compared to HC in serum. Gene sets were retrieved from MsigDB version 7.5 (Liberzon et al., 2015), pathway database source encompasses: KEGG (Kanehisa et al., 2023), REACTOME (Gillespie et al., 2022), BioCarta (Nishimura, 2001), PID (Schaefer et al., 2008). Only curated pathways from MsigDB were used (C2), those archived by MsigDB were excluded. In total, 22 gene sets satisfying the former criteria were considered.

### 2.2 Methods

#### 2.2.1 singIST method

We start by defining the three inputs of singIST: superpathways, human scRNA-seq, and disease model scRNA-seq Fold Changes (FC). First, we define the concept of a superpathway 𝒫^*p*^ as a set containing cell types and genes. We name *C*^1^, …, *C*^*b*^, …, *C*^*B*^ as the cell types of interest, previously identified and annotated. For each superpathway 𝒫^*p*^, there is a gene set 𝒢_*p*_ extracted from a pathway of interest *p*, from which gene subsets are derived for the cell types 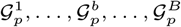 where 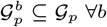. Each superpathway is formally defined as 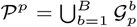, with the complete set of superpathways 𝒫 = {𝒫^1^, …, 𝒫^*p*^, …, 𝒫^*P*^} representing all pathways under evaluation. singIST method runs independently for each of the superpathways; hence, without loss of generality, from now on we fix a superpathway 𝒫^*p*^.

Second, we structure the human scRNA-seq data according to the superpathway. Let 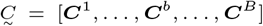 be the block of matrices containing the pseudobulk log-normalized expression for each cell type. Each matrix is defined element-wise 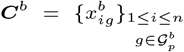, where 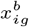 is the pseudobulk gene expression of human sample *i* for gene *g* in cell type *b*. We assume that human samples belong to different experimental groups *k* = 1, …, *K*, from which we define the *target class k* = *k*_1_ as the human experimental group that the disease model is intended to mimick (i.e disease, treated, etc.), and the *base class k* = *k*_0_ as the human experimental group that should differentiate from the *target class* (i.e healthy control, untreated, etc.). Let ***Y*** be the response matrix denoting human sample class, defined element-wise 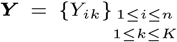, where elements *Y*_*ik*_ satisfy (1):

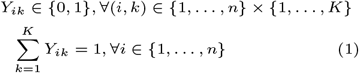

Third, we assume there are *l* = 1, …, *L* disease models to be assesed against human scRNA-seq data for each superpathway. Since singIST runs independently for each disease model, without loss of generality we fix a disease model *l*. We structure the disease model scRNA-seq FC as blocks of vectors. Let 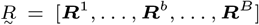 be the block of vector containing the FC between *target class* and *base class* of disease model samples. Each vector is defined element-wise 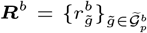, where 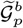 denotes the human gene subset 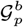 with its equivalent gene organism symbols for the disease model. The FC 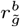 are computed through Eq (2).

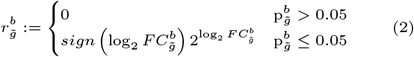

Where 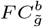 and 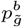 are the logFC of *target class* versus *base class* disease model samples and its adjusted p-value, respectively, both obtained from FindMarkers of Seurat.

We choose adaptive sparse multi-block partial least square discriminant analysis (asmbPLS-DA) as the basic model for our method, since it discriminates between multiple disease outcome groups and selects features on high-dimensional omics data using a multi-block data structure (Zhang and Datta, 2023). The detailed introduction of asmbPLS-DA is shown in Supplementary Material S1.

asmbPLS-DA is trained on human data, with ***Y*** as the response and 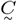 as the block-variable, always centered and scaled, with *k*_0_ being the reference class. Therefore, for each PLS component *j* = 1, …, *J*, we obtain estimates of response *q*, cell *ω*^*super*^ and gene *ω*^*b*^ weights indicating their relevance on discriminating between classes. Figure 1 depicts the trained model. Once the asmbPLS-DA model is trained, we can then predict the response ***Y*** for the original predictor data 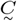 and new samples. Concretely, with the original data 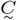 and estimated cell and gene weights, the scores *t*^*super*^ and *t*^*b*^ are calculated at cell and gene level, respectively, with which we predict the fitted model Eq (3).

**Fig. 1.**
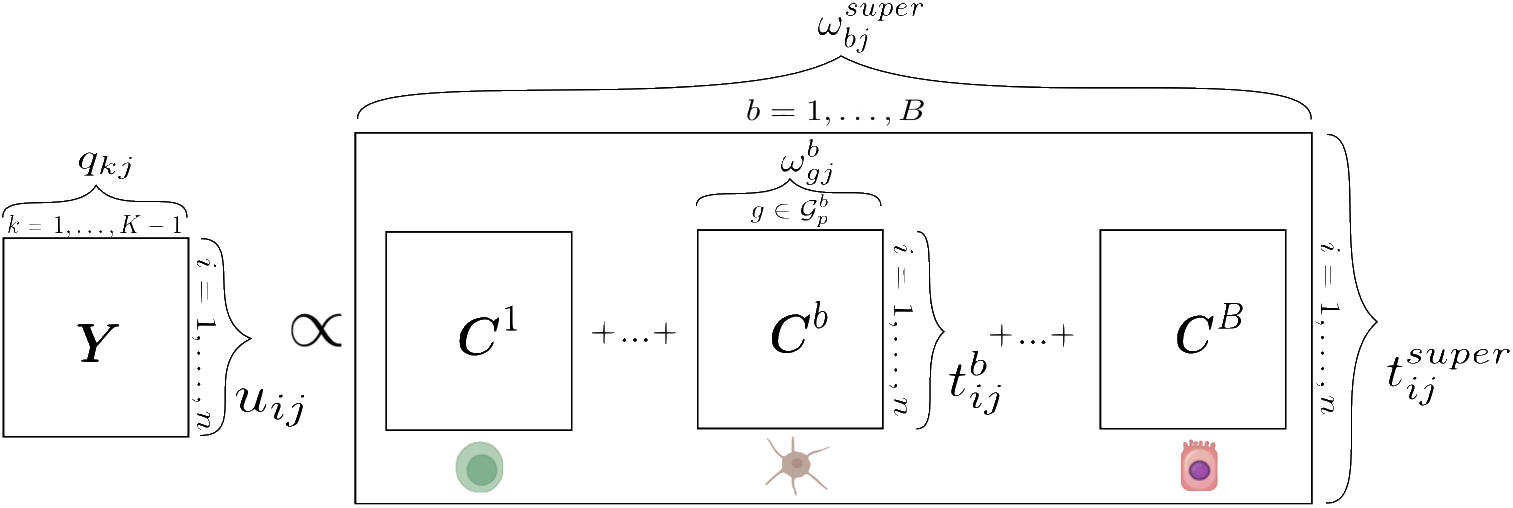
Representation of asmbPLS-DA for scRNA-seq readouts. The response matrix ***Y*** contains samples as rows and classes as columns, one-hot encoded. Predictor blocks ***C***^*b*^ are defined by cell types, with columns representing genes and rows representing samples. Each element within these predictor blocks is the pseudobulk of gene expression values. The figure displays loadings for the response matrix ***q***_*kj*_, predictor blocks 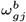 and the predictor superblock 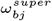, as well as scores for the response matrix ***u***_*ij*_, predictor blocks 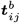, and the superblock 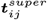.

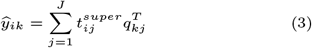

For a generic human sample and class, we name the continuous response prediction *ŷ*_*ik*_ as the superpathway’s score.

We define in Eq (4) the *superpathway reference recapitulation* as the difference in the median superpathway’s score between the *target class* and *base class* samples.

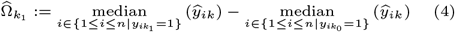

In Supplementary Material S1 we show that Eq (3) can be further developed into Eq (5) and Eq (7).

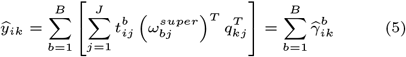

Where 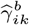 is the contribution of cell type *b* to the superpathway’s score of class *k*. With the former contributions, we characterize Eq (6) as the *cell type b reference recapitulation* defined by the observed difference in median cell contribution *b* to superpathway’s score between the *target class* and *base class*.

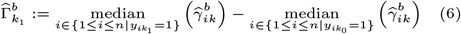

Similarly, we can extract such contributions at the gene level as shown in Eq (7).

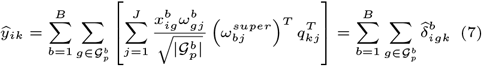

Where 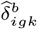 is the gene *g* contribution to cell type *b*. Note that 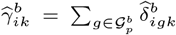, hence the cell type contributions to the superpathway’s score are the sum of all gene contributions within the cell type.

With the reference recapitulations, we have built a set of metrics based on the initial superpathway 𝒫^*p*^ that inform us on the similarity, at varying granularity levels, between *target class* and *base class* human samples.

Our aim now is to assess how similar the disease model is to the *target class* human single-cell gene expression. However, a direct comparison is not possible. For this reason, we define the *singIST treated samples* as the single-cell human gene expression samples we would have observed if human scRNA-seq behaved like the changes observed in the disease model. This is an assumption that IST and FIT follow to model human gene expression as a function of disease model changes measured through FC. With the disease model FC we derive the *singIST treated samples* by using Eq (8) in human samples belonging to the *base class*.

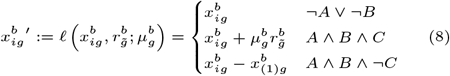

Where 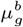 is the gene centroid computed by asmbPLS-DA, an intuition of the *Biological link function* 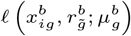 is shown in Supplementary Material S1. Three biologically plausible scenarios define the transformation in Eq (8); (¬*A* ∨ ¬*B*) the case where either cell type *b* does not exist in the disease model (not A) or disease model does not have a one-to-one ortholog of human gene *g* (not B); (*A* ∧ *B* ∧ *C*) the case where both cell type *b* and a one-to-one ortholog gene of *g* exist in disease model, and the translation of FC does not produce a negative expression 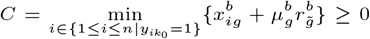; (*A* ∧ *B* ∧ ¬*C*) the case when both cell type *b* and one-to-one ortholog gene of *g* exist in the disease model, and the translation of FC does produce a negative expression any human sample (not C).

With the new predictor block 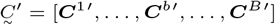 we use Eq (3) to predict the continuous response **Ŷ**^**′**^. Equally as before, we can define the *predicted recapitulations* of the disease model.

The *superpathway predicted recapitulation*.

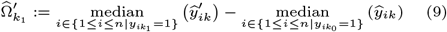

Likewise, the *cell type b predicted recapitulation*.

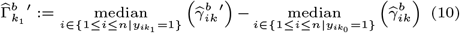

To ease interpretation on the similarity between human and disease model, one can express the predicted recapitulation as a fraction of the reference recapitulation, at both granularity levels: superpathway 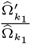 and cell type 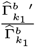. In Supplementary Material S1 we prove the former fractions depend on: direction and magnitude of FC (2), orthology 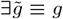, cell type existence in disease model, and human gene *ω*^*b*^ and cell type weights *ω*^*super*^.

The *superpathway predicted recapitulation* as a fraction of the *superpathway reference recapitulation*.

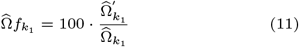

*Cell type b predicted recapitulation* as a fraction of *cell type b reference recapitulation*.

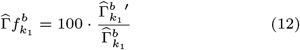

To allow for downstream analysis on what drives cell type recapitulation, we define the contribution of gene 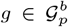 on recapitulation (12).

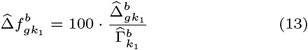

Where 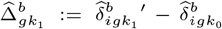, which is constant for all samples *i*, this fact is proven in Supplementary Material S1. Interpretation of (13) is the contribution of gene *g* toward *cell reference recapitulation* 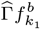, this is due to the relationship between (12) and (13) as 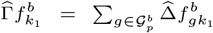 (proof in Supplementary Material S1).

A graphical summary of singIST procedure is shown in Fig 2.

**Fig. 2.**
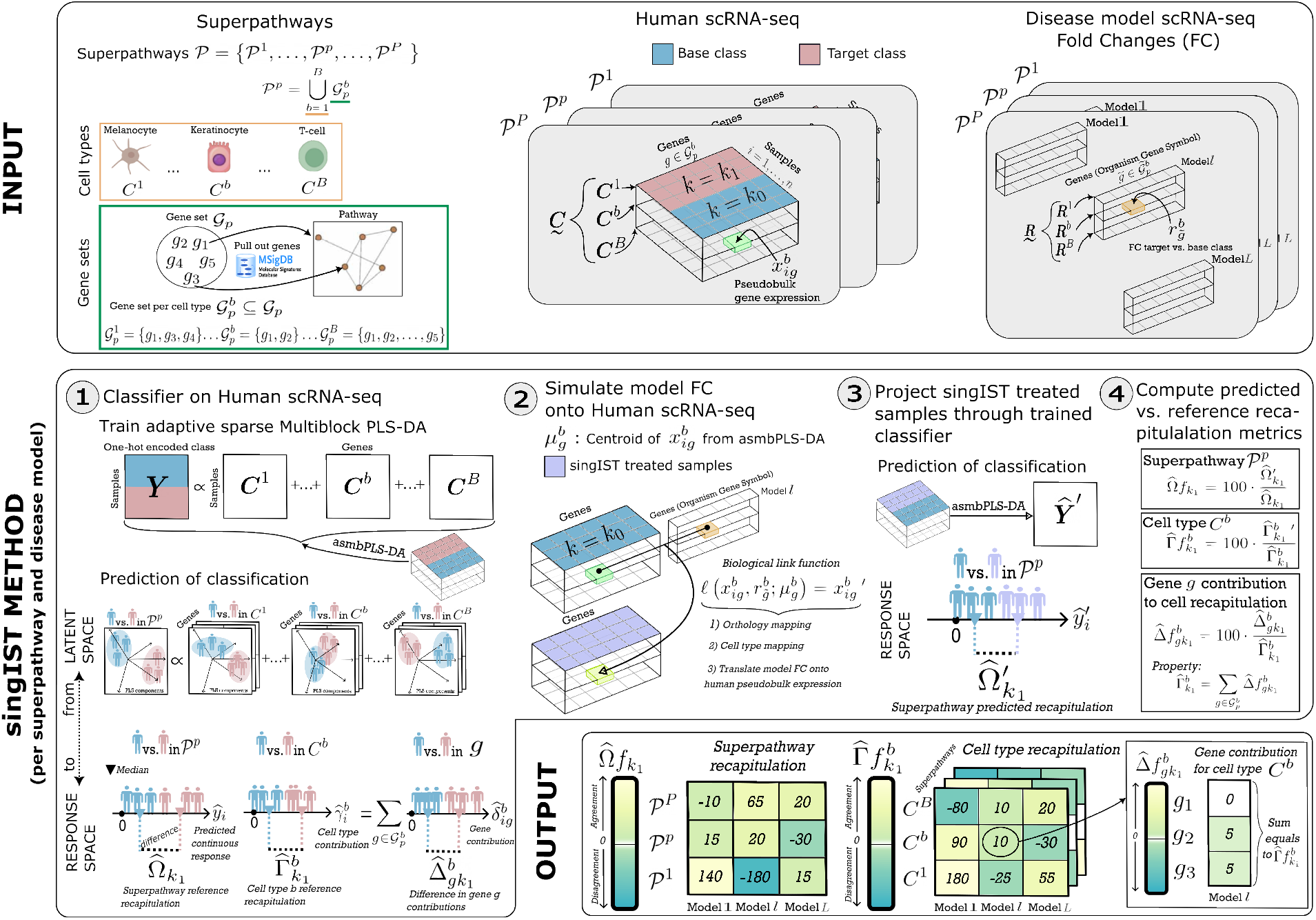
INPUT) First, definition of a superpathway (𝒫^*p*^) as a set containing cell types and genes. For each 𝒫^*p*^, there is a gene set 𝒢_*p*_ from which gene subsets are derived for cell types 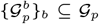. Second, for each 𝒫^*p*^ human scRNA-seq data is organized into matrix layers. Target class (*k* = *k*_1_) is the human experimental group that the disease model aims to mimick (i.e disease, treated), while base class (*k* = *k*_0_) is such that should differentiate from target class (i.e healthy control, untreated). Third, for each 𝒫^*p*^ disease models scRNA-seq FC are structured into vector layers. **singIST METHOD)** The method is organized into four steps, which runs independently for each 𝒫^*p*^ and disease model. **Step 1)** *Objective:* Quantify differences between target and base classes human samples at various levels of granularity (superpathway, cell type, and gene) using asmbPLS-DA. *Input:* A 𝒫^*p*^ and human scRNA-seq data. *Output:* Optimal asmbPLS-DA. From such, we derive cell type contributions 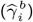 and gene contribution 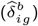. With the contributions we compute similarity measures at the superpathway 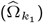 and the cell type levels 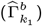. **Step 2)** *Objective:* biologically unify the human data with the disease model data for subsequent comparison. *Input:* Human scRNA-seq base class samples and disease model scRNA-seq FC data. *Output:* Human scRNA-seq gene expression observed when disease model FC are applied, we call them *singIST treated samples*. The former is achieved in the *Biological link function*, which performs steps; one-to-one orthologous mapping; cell type alignment; translate FC to 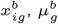 is a scaling factor. **Step 3)** *Objective:* Compute metrics of output from Step 1 between singIST treated samples and human scRNA-seq base class. *Input:* singIST treated samples, Human scRNA-seq base class samples and optimal asmbPLS-DA. *Output:* Pathway predicted recapitulation 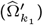, Cell type *b* predicted recapitulation 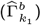 and predicted gene contributions 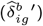. **Step 4)** *Objective:* Compute similarity metrics between human and disease model. *Input:* From step 1; 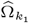 and 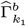, and 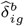. From step 3; 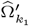 and 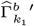, and 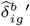. *Output:* Predicted recapitulations as a fraction of the reference recapitulations 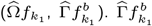 is explained by contributing genes 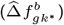 providing interpretation on which genes drive the cell type recapitulation. **OUTPUT)** 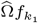 and 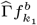 are displayed. Positive values show agreement in gene expression change between disease model and humans, negative show opposed one. Each 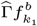 equals to the sum of its gene contributions 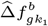.

#### 2.2.2 Interpretation of recapitulation measures

Recapitulation measures have a related interpretation to that of IST (Picart-Armada et al., 2024). A recapitulation of 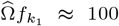 would imply that the median of the predicted superpathway’s score of singIST treated samples corresponds to that of the *target class*, hence human data show similarity to disease model. If cell recapitulation 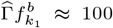 then the median of the predicted cell contribution to superpathway’s score of samples singIST treated samples corresponds to that of the *target class*, and human data show similarity to disease model for cell type *b*. Gene contribution 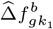 to cell type recapitulation *b* provides with the magnitude of change and direction such gene has contributed to 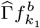. This contributions will vary greatly depending on the case; positive gene contributions arise from large 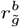 in disease model and agreement in direction of change in the asmbPLS-DA model in human data; negative contributions arise from opposed directions of change between 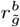 and that estimated in the asmbPLS-DA with human data. On the other hand, if a cell type is not present in disease model its cell recapitulation 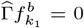, and consequently all its gene contributions are null. Similarly, if gene a does not have a one-to-one disease model ortholog 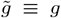 then 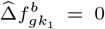. Genes having no relevance in asmbPLS-DA and/or null differences in disease model would have a recapitulation 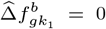. Details on these facts are shown in Supplementary Material S1.

### 2.3 Validation methodology

#### 2.3.1 Validity test of the optimal asmbPLS-DA

Once the optimal model is selected, the validity of such for classifying between *k*_1_ and *k*_0_ is checked by a permutation test (Brandolini-Bunlon et al., 2019) adapted to small sample size and asmbPLS-DA. A null model distribution 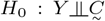 is generated by permuting Y, noted as *σ*(*Y*), and setting the number of permutations. For each *σ*(*Y*) an asmbPLS-DA model is fitted with *J*^*^ and 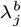 as the optimal number of PLS components and quantile combination for each block and PLS component, respectively, and randomly taking one sample out to avoid overfitting. With the permuted model, *Y* is predicted for all samples under analysis, such prediction is compared against the true *Y* by *F*_1_ score. The rationale behind randomly permuting each *Y* element is that its original relationship of the model is disrupted while the dependence structure of 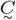 is preserved (Winkler et al., 2015), thus providing a control of a false positive model. If the optimal model is actually significant it is expected that error measures increase substantially when permuting.

To this end, the *F*_1_ LOOCV error of optimal model is compared against the *CI*_(1−*α*)_ of null distribution of *F*_1_ score, where *α* = 0.05 is the confidence treshold and the quantile serves as the p-value which is adjusted for multiple comparison by Benjamini-Hochberg (Benjamini and Hochberg, 1995), FDR is set to 0.1.

#### 2.5.2 Parameters variabilities and significance of the optimal asmbPLS-DA model

Cell type and gene importance, within a cell type *b*, may be assessed by considering the weighted average of its estimated coefficient, by taking as weights the relative importance of each PLS component 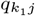 (Bougeard et al., 2011). To this end, we define the Cell Importance Projection (CIP) for cell type *b*:

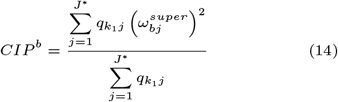

Similarly we define the Gene Importance Projection (GIP) for gene *g* within cell type *b*:

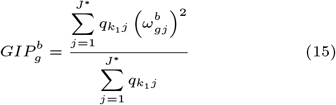

Both indices verify 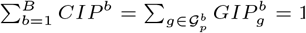, this is proven in Supplementary Material S1. The direction of *CIP*^*b*^ may be assesed by 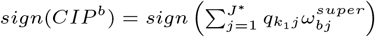, and equivalently for 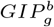. Note that *CIP*^*b*^ distribution is nested to the already estimated 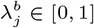, the blocks with only small number of relevant genes will assign higher 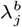 values, being a cell type 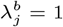 a cell type that does not contain any relevant information in classifying between *k*_1_ and *k*_0_.

The 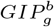 distribution of a gene that is significant is likely to substantially differ from its associated null *H*_0_ distribution. A null distribution of 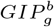 of the form 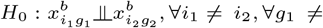, and *b* ∈ {1, …, *B*} is generated by permuting all samples and genes within block. Note that permuting between block would not satisfy exchangeability assumption as 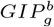 distribution is dependent on *λ*^*b*^. The median 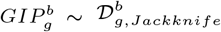 of distribution of the optimal model is compared against the null distribution 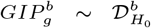 by a Mann-Whitney U test, where the alternative hypothesis being the median 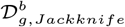 greater than median of 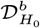. P-value is adjusted by Bonferroni correction with a lower bound of expected number of true null hypothesis for each cell type 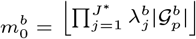, the rationale is provided in Supplementary Material S1.

## 3. Results

All tables and figures show only superpathways under discussion, one can find tables and figures with the full 22 superpathways in Supplementary Material S3. Table 2 provides a summary of characteristics of the optimal trained models, gene set size per cell type 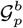 was set equal to 𝒢_*p*_ for all pathways. The gene set sizes of pathways are highly variable, ranging from a minimum of 15 genes for CD40/CD40L signaling [PID] to a maximum of 701 genes for Cytokine signaling to the Immune system [REACTOME]. All optimized asmbPLS-DA demonstrated statistical significance at a false discovery rate (FDR) treshold of 0.1. The optimal quantile sparsity values for Dendritic Cells in Th1/Th2 Development [BIOCARTA] show that T-cell (TC) are the mayor source of information with *λ*^*b*^ = 0.35 and 4 statistically significant genes, followed by Melanocytes (MC) with *λ*^*b*^ = 0.45 and 4 statistically significant genes, and Dendritic Cell (DC) *λ*^*b*^ = 0.55 and 3 statistically significant genes. On the contrary, both Keratinocyte (KC) and Langerhans Cell (LC) show the least importance with *λ*^*b*^ = 0.95 and 1 statistically significant gene. For JAK-STAT signaling pathway [KEGG], all cell types show the same amount of relevant information with *λ*^*b*^ = 0.05, suggesting an active role of all cell types in activating this pathway, with the number of statistically significant genes ranging from a minimum of 32 for TC and LC, and a maximum of 41 genes for MC. In Chemokine receptors bind chemokines [REACTOME], KC and DC were the most informative cell types with both showing a *λ*^*b*^ = 0.05 and 20 and 13 statistically significant genes, respectively. Lastly, Antigen Presenting Cells (APCs) were the most informative in Cytokine-Cytokine receptor interaction [KEGG] with 50 statistically significant genes for LC and 31 for DC.

**Table 1.**
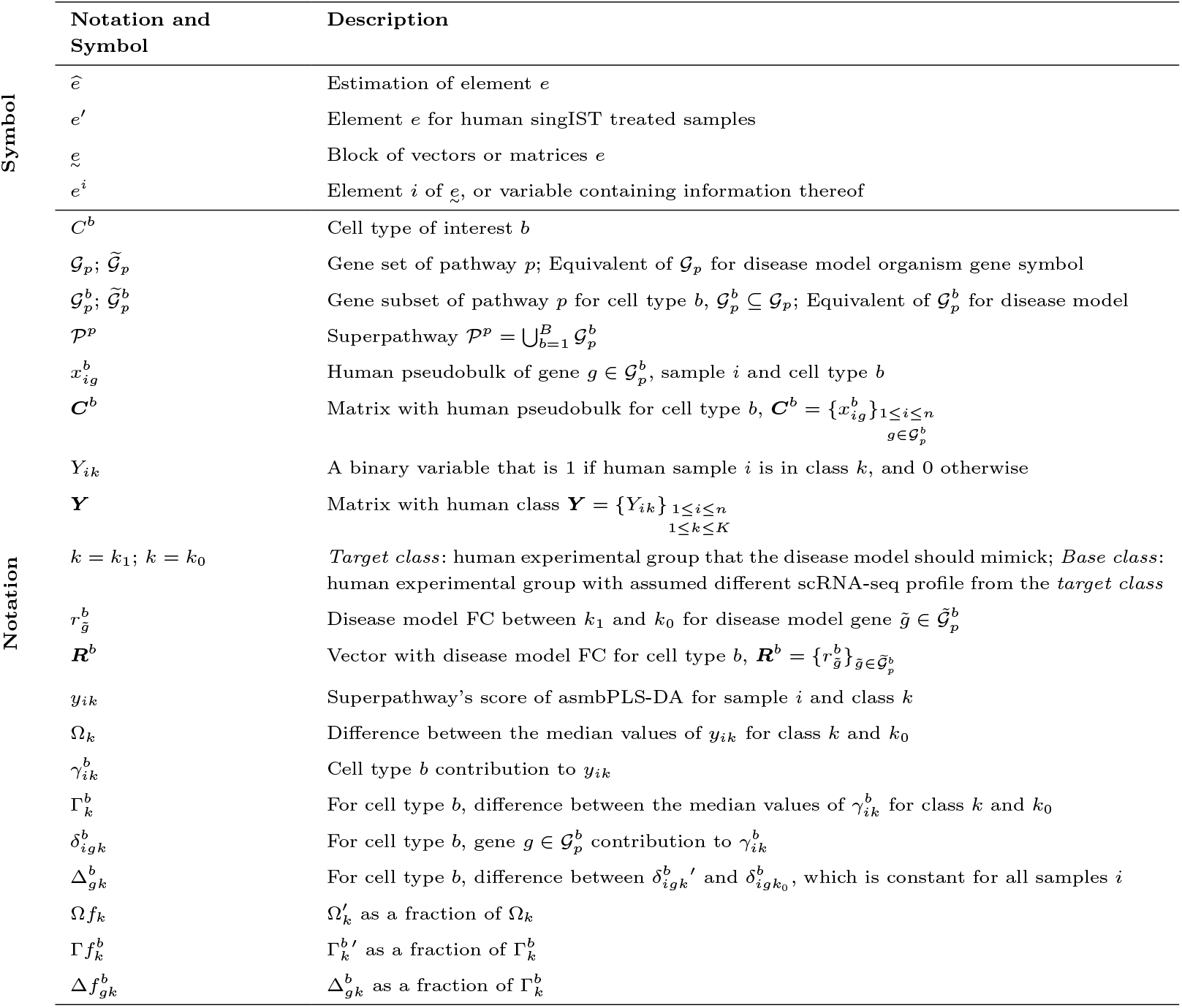
Summary table of main symbols and notations defined in singIST method.

**Table 2.**
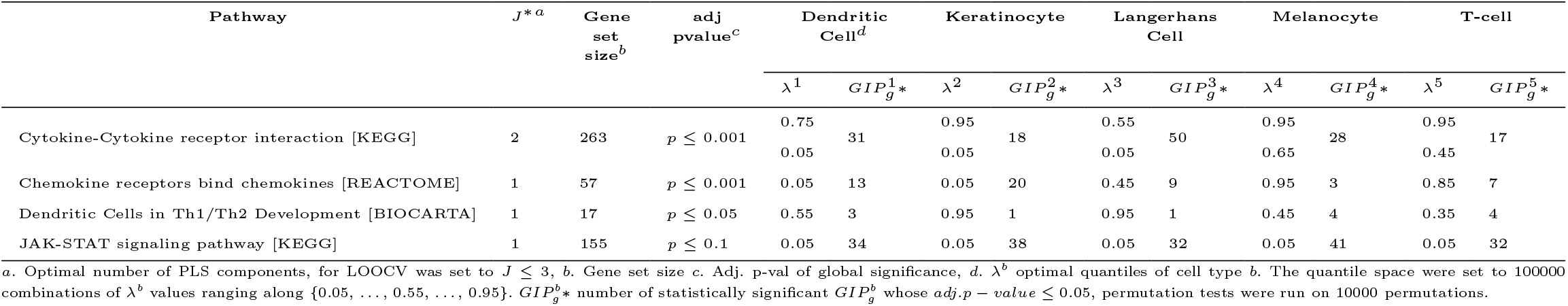
Characteristics of fitted optimal asmbPLS-DA for all superpathways.

To offer an overview of the most predictive genes for classifying between AD lesional and HC, Table 3 lists the five statistically significant genes with the highest 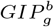 values, along with their corresponding directional references from the literature. In Dendritic Cells in Th1/Th2 Development, it is genes within TC that drive the prediction, concretely, IL13 shows the strongest upregulation, along with IL5 and CSF2/GM-CSF, while TLR7 shows a downregulation. ANP/CD13 shows a strong upregulation in both APCs, while for MC it is an upregulation of IL10 that drives the prediction, as well as the upregulation of antigen coding genes: ITGAX/CD11c, CD7 and CD33. All cell types in JAK-STAT present a similar importance, in line with the *λ*^*b*^ obtained, however top five genes within each cell type differ. In TC higher expression in AD samples compared to HC of interleukins dominate IL13, IL26, IL2RA, IL7, as well as IFNGR1. MC from AD samples show a simultaneous upregulation of CCND3 and CCND1, and so do receptors IFNGR2 and IL10RA. KC are characterized by overexpression of IL15 and its receptor IL15RA, on the contrary we observe a downregulation of SPRIY1. Further, IL23A and IFNL1/IL-29 are downregulated in DC, while SPRED1, SOCS1 and OSM are upregulated. For LC a downregulation of genes IL22RA2, JAK1 and CCND1 is estimated, on the contrary there is a downregulation of CCND2 and STAT6. APCs are the mayor drivers of Cytokine-Cytokine receptor interaction pathway predictivity. Precisely, for DC chemokine receptor and ligands CCR6, CCL5/RANTES and CCL3L1 were all supressed in AD, and Herpesvirus entry mediator gene TNFRSF14 was upregulated. IL23A is inhibited in both LC and DC. Chemokine receptors bind chemokines pathway are fueled by KC, DC and LC. Common chemokine AD markers are observed; CCL20/LARC, CCL7/MCP1 and CCR2 upregulatione in KC; upregulated CCL17/TARC, CRR1 and CXCR4 in LC; CCL13 in DC.

**Table 3.**
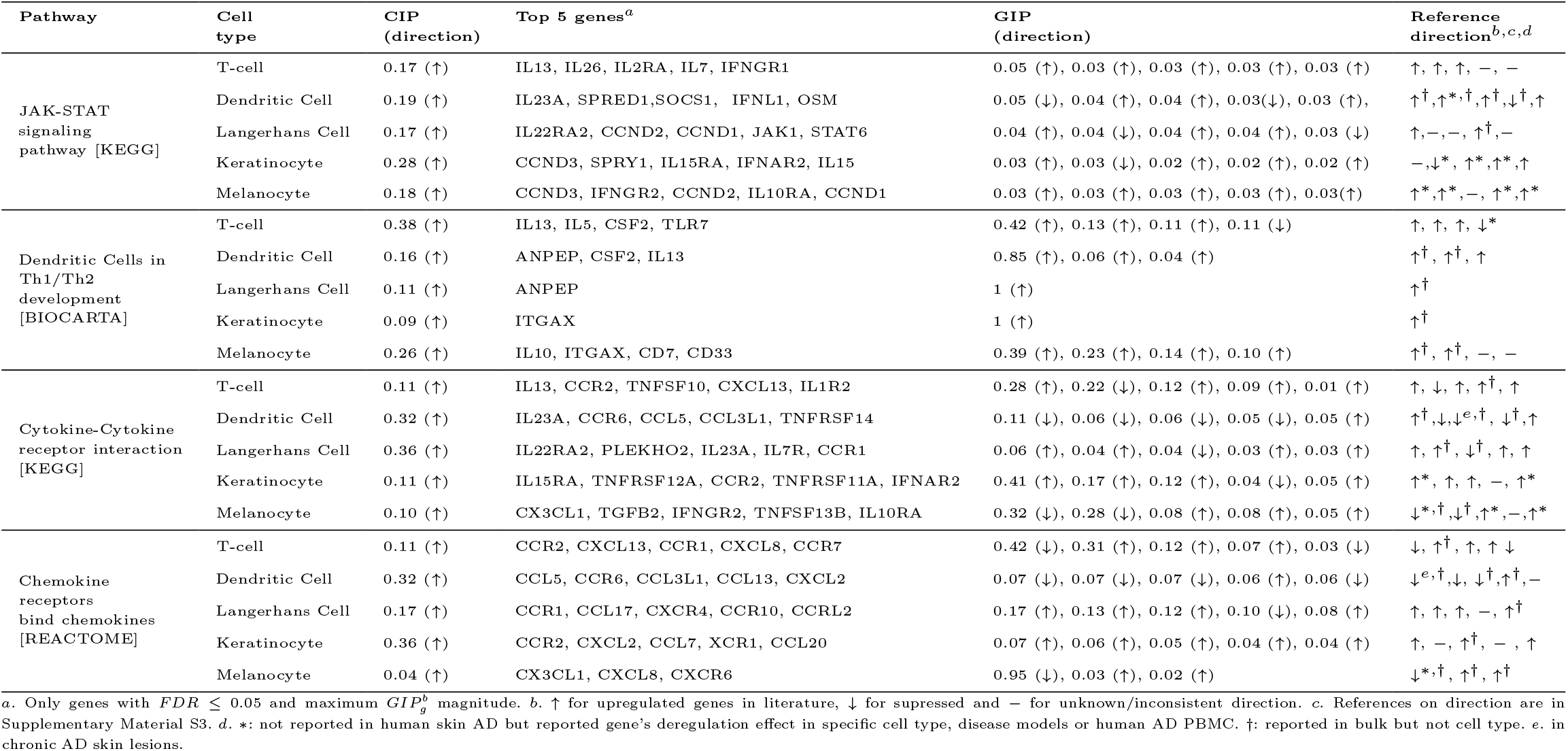
Top five statistically significant genes ordered by 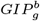 magnitude.

One-to-one orthology of each disease model against humans was assessed, likewise their superpathway recapitulations from singIST procedure are shown in Figure 3. All disease models exhibit a similar level of observed one-to-one orthology, with OVA typically displaying fewer sequenced genes per pathway. However, orthology coverage varies significantly by pathway. Notably, the Asthma and Chemokine receptors bind chemokines pathways show 60% orthology coverage, while the IL12 signaling mediated by STAT4 pathway exhibits 100% coverage. Superpathway recapitulations for Dendritic Cells in Th1/Th2 Development show overall agreement in direction with human AD for OXA (96.6%), as the highest recapitulation, followed by IMQ (41.4%), and OVA (0%) showing no recapitulation at all. Cell type recapitulations are displayed in Figure 4, which show high TC recapitulation for IMQ (136%) and no recapitulation for OVA (0%) and OXA (1.3%). Concretely, OXA is driven by LC (549.7%) and DC (204.3%) recapitulations. OVA has 0% recapitulation for LC across all pathways, since the LC cell type had fewer than 100 cells in OVA, they were removed. MC have 0% recapitulation across all disease models and pathways. Figure 5 illustrates the gene contribution to cell type recapitulation. TC recapitulation of IMQ come from CSF2/GM-CSF (127.8%) and IL5 (45.7%), while TLR7 (−39%) shows disagreement with human condition, since its FC is 6.1 while TLR7 is suppressed in human AD. OXA recapitulation is solely driven by ANPEP with strong FC in LC (13.3) and DC (2.8), which aligns with the observed upregulation of ANPEP in human APCs. OVA stands out for having no DEGs in this pathway, resulting in 0% recapitulation. Interestingly, strongest TC marker IL13 in AD is not DEG by any disease model. In the JAK-STAT signaling superpathway recapitulations, OXA (84.2%) performs the highest while IMQ (13.9%) and OVA (1.5%) show low recapitulations. Precisely, OXA agrees in direction with moderate recapitulations for all cell types, strongest contributing genes are IL21R (−31.3%) in DC, OSM (58.1%; 107.6%; 61.2%) across APCs and KC, and all STAT1/2/3/5 in TC. Gene contributions in TC for IMQ generally agree with OXA with disagreement in direction and/or magnitude on: CSF2 (15.2%), CSF3R (−1.7%), IL23R (−1%), IL2RG (−2%) and JAK1 (−5.1%). Cytokine-Cytokine receptor interaction pathway show no superpathway recapitulation at all for IMQ (−1.3%) and OVA (0.6%), while OXA (376%) a very extreme recapitulation. For IMQ and OVA, cell type recapitulation is very heterogeneous with some agreeing on direction (TC, LC) and others disagreeing (KC, DC), which overall drives the superpathway recapitulation to almost zero. The same holds for OVA. OXA extreme recapitulation in LC (792.8%) comes from INHBA (687.1%) with an extreme FC of 627.2, while disagreement in KC (−80%) are mainly from CXCR2 (−44.4%), OSM (−21.8%), CRR5 (−8.8%). Superpathway recapitulations of Chemokine receptors bind chemokines depict high differences between the murine models, high recapitulation for OVA (105.2%), disagreement in direction for IMQ (−25.5%) and OXA (−163.2%). OVA recapitulation comes purely from KC (284.4%) with CXCL6 (280%) being the mayor source of contribution due to its extreme FC (499.6) and agreement with human direction. On the contrary, OXA and IMQ have higher variability between cell type recapitulations, while they agree on LC recapitulation direction, they disagree on DC and KC with IMQ (−66.6%; 6.6%) and OXA (77.8%;-610.9%). The source of differences for DC are CCL5 (IMQ −91.5%, OXA 0%), and for KC is CXCR2 (IMQ 0%; OXA −483.3%). While both IMQ and OXA show opposite directions to that of humans in TC, both mainly due to disagreement in CCL5 and CCR2.

**Fig. 3.**
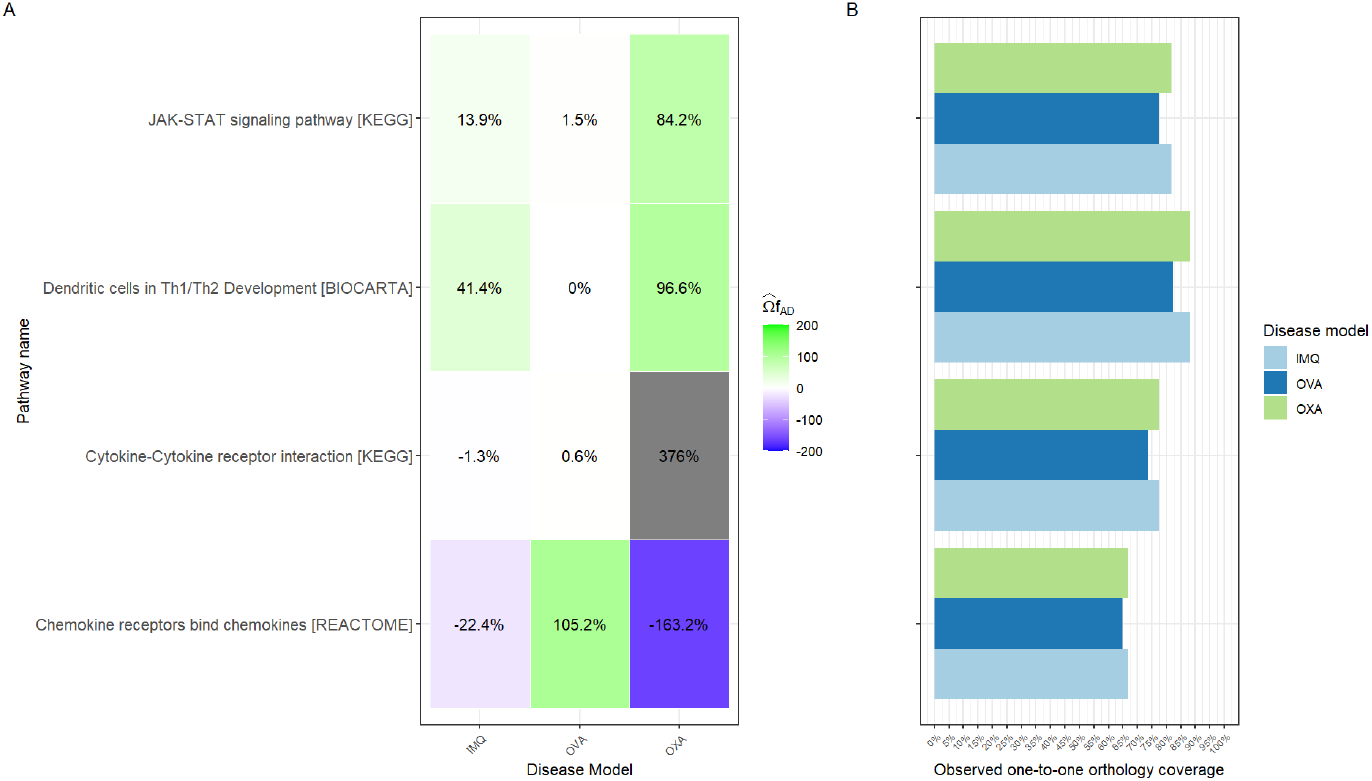
Superpathway recapitulation and observed one-to-one orthology of AD disease models. **A)** Superpathway predicted recapitulation as a fraction of the superpathway reference recapitulation for IMQ, OXA and OVA across all pathways under study. Negative recapitulations refer to opposed directions with human observed condition, while positive recapitulations define agreement in direction. **B)** Observed one-to-one orthology coverage refers to number of observed and one-to-one ortholog genes in disease model as a fraction of pathway gene set size. Despite all disease models belong to the same organism *mus musculus* their differences in observed orthology one-to-one coverage come from sequenced reads.

**Fig. 4.**
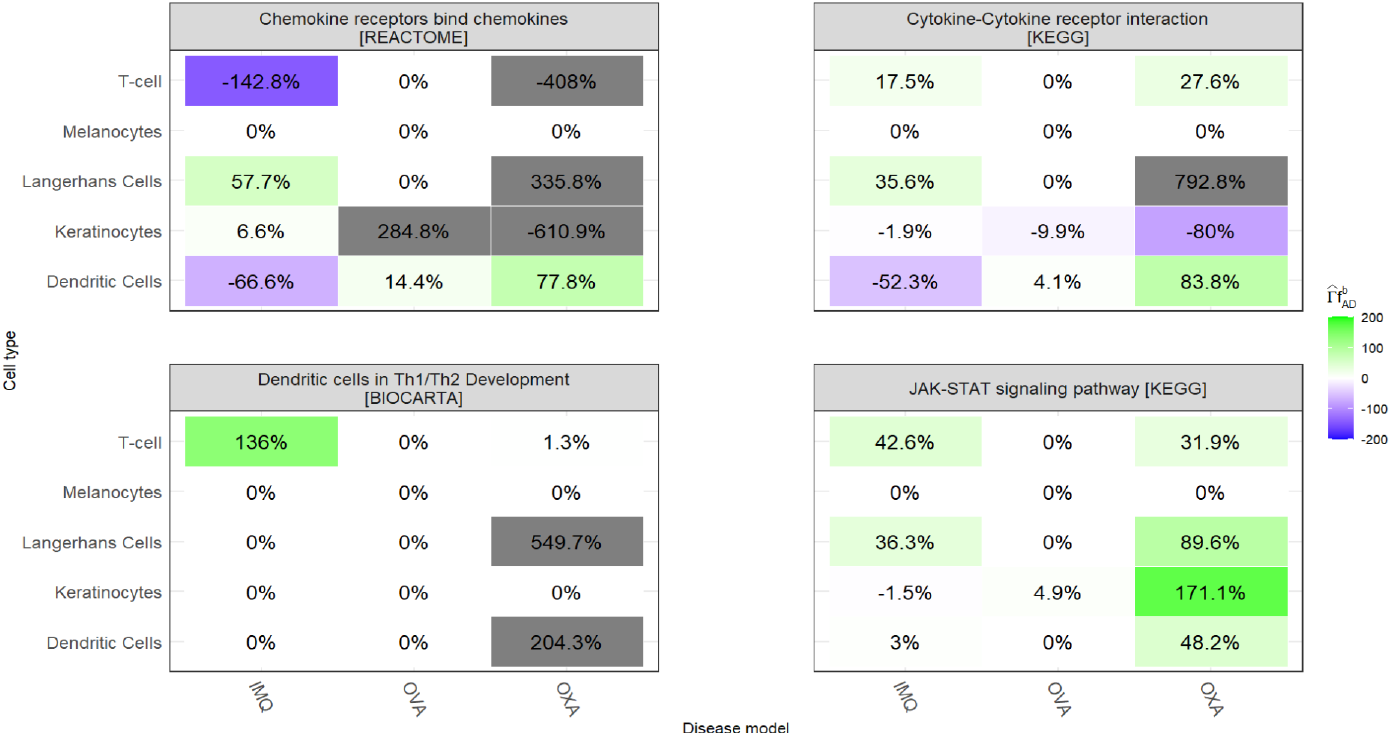
Cell type recapitulation for all AD disease models and superpathways under analysis.

**Fig. 5.**
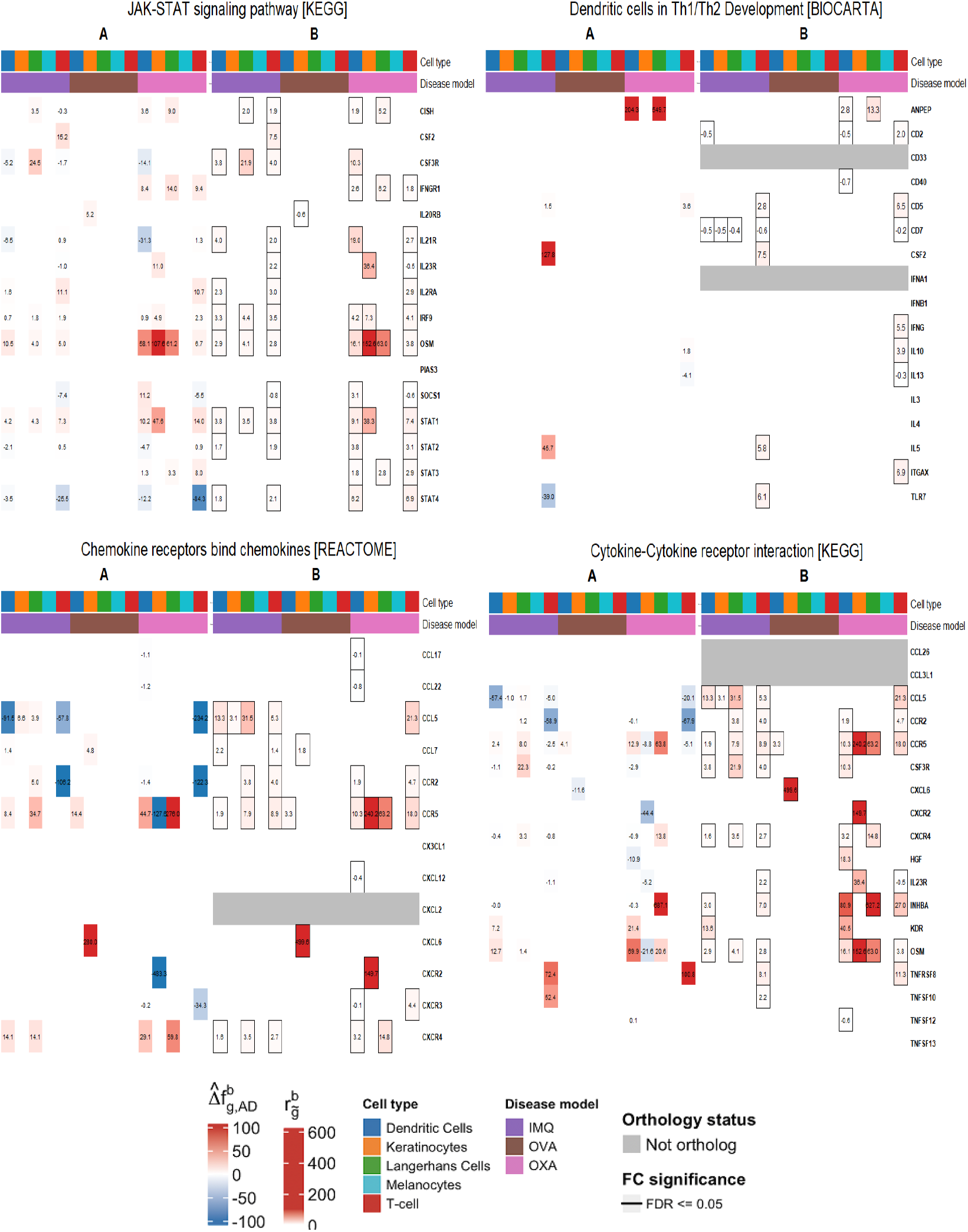
Gene contribution and disease model estimated 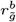. **A)** Gene contribution to cell type recapitulation by disease model. If gene set size of pathway is greater than 50, only the top 5 contributing genes, for each cell type, were displayed. Blank gene contributions correspond to 0 values. **B)** Computed 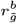 by disease model. Grey FC refer to genes without one-to-one ortholog and/or not sequenced in disease model. Framed FC refer to statistically significant *FDR* ≤ 0.05 genes, as per FindMarkers. Blank FC correspond to 0 values.

## 4. Discussion

Atopic Dermatitis (AD) represents a chronic skin-immune-mediated inflammatory disease (IMID), characterized by dysregulated T-cell mediated inflammation and keratinocyte differentiation (Tsoi et al., 2019). We put a special focus in the discussion on pathways proven to be causal drivers of AD pathogenesis or related to its clinical severity; JAK-STAT signaling pathway, Dendritic Cells in regulating Th1/Th2 development, Cytokine-Cytokine receptor interaction, and Chemokine Signaling pathway. As anticipated, distinct cell types predominantly activate each pathway: TC and DC in Dendritic Cells in Th1/Th2 Development, KC and DC in Chemokine receptors bind chemokines, LC and DC in Cytokine-Cytokine receptor interaction. Conversely, in the JAK-STAT signaling pathway, all cell types play a significant role in its activation. The JAK-STAT pathway is involved in a variety of biological functions beyond immunity, including but not limited to: cell division, cell death, and tumor formation. AD is characterized by intense itching and scratching, which leads to eczematous lesions. These lesions damage MC and KC which in turn activate the JAK-STAT signaling pathway, promoting cell-cycle and inflammation. It is reported that simultaneous activation of CCND3/Cyclin-D3 and CCND1/Cyclin-D1 in melanocytic skin lesions promotes cell-cycle regulation (Alekseenko et al., 2010). Additionally, MYC is upregulated and involved in this function. Furthermore, unique genes in the PIK3T-AKT subpathway, such as PIK3CD and PIK3CB, which are involved in cell-cycle and cell survival functions, are also upregulated in the JAK-STAT pathway. Similarly, upregulation in KC of IL15RA/IL15R*α* and IL15 in skin lesions has been associated to inflammatory processes. Additionally, the activation of the JAK-STAT pathway by TC, LC and DC is primarily related to immune functions.

It was interesting, though not surprising, to observe the low number of DEGs in OVA for the pathways under study. This finding aligns with the low DEGs observed in TC and DC clusters reported in Leyva-Castillo et al. (2022), as well as bulk-RNA studies showing a low number of DEG in characteristic pathways of AD (Ewald et al., 2017). The absence of MC across all disease models is expected, as the interfollicular epidermis of mouse pelage skin, particularly in the ear where biopsies are collected, entirely lacks functional, pigment-producing MC (Michalak-Mićka et al., 2022).

Dendritic Cells in Th1/Th2 Development pathway varies significantly across different disease models. In the IMQ model, there is a notable upregulation of TLR7 (FC 6.1), whereas this gene is suppressed in human AD, leading to a negative gene contribution of −39%. Although TLR7 has not been extensively studied in AD, its role in regulating TC differentiation is well-documented. Suppression of TLR7 in TC has been shown to produce a skewed Th2 immune response (Jeisy-Scott et al., 2011), consistent with the immune response observed in AD. Conversely, upregulation of TLR7 in murine models leads to the differentiation of TC towards Th17 and/or Th1 (Ye et al., 2017). In the IMQ model, upregulation of TLR7 is expected for two reasons; first, the mechanism of action of Imiquimod cream is as a TLR7 agonist; second, IMQ is characterized by a Th17-driven immune response. However, in human AD, suppression of TLR7 would be anticipated due to its Th2-skewed response. Furthermore, it is known that none of these disease models accurately mimic the mechanisms that mediate expression of IL13/IL4 genes. While the IMQ model is characterized by an increase in IL17/IL22 and not IL13/IL4, typical of psoriatic lesions, OVA and OXA models produce IL13/IL4 through infiltrating basophils (myeloid cells) rather than Th2 cells (Leyva-Castillo et al., 2022; Liu et al., 2020), contrary to previous thought, leading to contribution of such genes of 0%, or even negative for IL13 in OXA (−4.1%).

In the JAK-STAT pathway, OXA demonstrated the best superpathway recapitulation at 84.2% with consistency in agreement across all cell types, while OVA performed the worst at 1.5%. OXA is preferentially chosen as a preclinical in vivo model for testing JAK inhibitors in AD (Zhang et al., 2024). One reason for OVA’s poor performance is its low number of DEGs in the JAK-STAT pathway. Previous bulk-RNA comparisons of OVA and OXA in the JAK-STAT pathway highlight the superiority of OXA (Ewald et al., 2017), underscoring the deficiency of low DEGs in OVA. In TC of OXA, large negative contributions were observed for STAT4, and moderate negative ones for DC and TC in both IMQ and OXA. STAT4 levels are greatly suppressed during physiological Th2 development (Usui et al., 2003), as observed in human AD. However, both IMQ and OXA immune responses are not characterized by Th2-skewed response, hence expression of STAT4 in DC and TC is upregulated, with strong FC in opposite direction to that estimated in human AD. A similar pattern is observed in DC for IL21R both in OXA and IMQ, while IL21R is stablished as an upregulated cytokine in skin human and murine AD lesions for TC (Jin et al., 2009), less its known about its role in DC. Both OXA and IMQ show positive small contributions aligned with human AD in TC for IL21R (IMQ: 0.9%, OXA: 1.3%) and negative ones for DC (IMQ: −6.5%, OXA: −31.3%).

The pathways Chemokine receptors bind chemokines and Cytokine-Cytokine receptor interaction show overlapping genes with large contributions. The expression of many chemokines and cytokines are not stable throughout the lifespan of the lesion; instead, their levels fluctuate depending on the stage of it. For instance, CCL5 shows large negative contributions in both pathways across TC for both IMQ and OXA, and additionally DC for IMQ. Particularly, CCL5 expression in human skin AD lesions is upregulated in the acute phase but suppressed in chronic AD wounds (Tsoi et al., 2020), here we observe a downregulation of CCL5 in human AD. Despite the lack of clinical information on lesion stages in Bangert et al. (2021), since all patients were diagnosed with chronic AD during early childhood, it is likely that their collected lesions are chronic. However, collected skin lesions from murine models are acute, which explains the difference in direction to human AD across all disease models and cell types. In addition, it is known that IMQ broadly upregulates CCL5/CCL4, while this is clear in CCL5 since TC, DC, LC and KC, show contributions, CCL4 is not a one-to-one ortholog with humans which leads to null contributions across all cell types. On the contrary, we observe CCL5 expression in OXA is more localized to TC as shown in Liu et al. (2020). CCL5 receptor CCR5 is also highly upregulated across all disease models, while OXA showing stronger contributions across all cell types compared to IMQ in line with Liu et al. (2020).

While demonstrating significant capabilities, singIST presents several limitations, including dependence on pre-annotated cell types, the assumption of homogeneous effects when translating fold changes to human gene expression, and the requirement for well-defined human disease states (e.g., endotypes) prior to analysis. Additionally, extensions could be explored on differentiating changes due to cell type-specific gene expression and cell type proportions, between human classes. Further validation in additional disease contexts will solidify its utility in drug development and preclinical research.

## 6. Conclusions

Here we have developed singIST an extension of IST method for comparative single-cell transcriptomics, offering a novel, integrative framework to evaluate disease model similarity to human conditions at various biological levels. singIST provides explainable and quantitative insights into transcriptomic alignment. Its application to murine models of atopic dermatitis demonstrated the method’s ability to recover known findings and generate novel hypotheses.

## Supplementary data

Supplementary Data are available online.

## Author contribution statement

A.M conceptualized and formalized the method, implemented it and performed the data analysis, interpreted data results and drafted initial article for review. S.P conceptualized the method. S.P, A.P and F.F revised critically the work for important intellectual content and approved the final version to be published.

## Acknowledgements

This work was supported by the Spanish Ministry of Economy and Competitiveness (www.mineco.gob.es) PID2021-122952OB-I00, DPI2017-89827-R, Networking Biomedical Research Centre in the subject area of Bioengineering, Biomaterials and Nanomedicine (CIBER-BBN), initiatives of Instituto de Investigación Carlos III (ISCIII), and with the support of the Pla de Doctorats Industrials de la Secretaria d’Universitats i Recerca del Departament d’Empresa i Coneixement de la Generalitat de Catalunya. B2SLab is certified as 2017 SGR 952.

A.M would like to acknowledge; Dr. Sergio Oller-Moreno and Tomas Romero-Rodriguez for software support; Dr. Bruna Oriol-Tordera and Mercè Pont-Giralt for providing useful references in AD and murine models; Dr. Juan Luis-Trincado for scRNA-seq bioinformatics support; Dr. Estrella Lozoya-Toribio for her summary skills.

