## Supplementary Material S1 for "singIST: an integrative method for comparative single-cell transcriptomics between disease models and humans"

### Supplementary Material S1: Materials and methods

#### A. Introduction to asmbPLS-DA

Here we introduce adaptive sparse multiblock partial least square discriminant analysis (asmbPLS-DA) (Zhang and Datta, 2023), a multiblock data fusion method that accounts for sparsity criteria adaptive to each predictor block. asmbPLS-DA is based on sparse multiblock partial least squares discriminant analysis (smbPLS-DA) (Li et al., 2012).

Let $\boldsymbol{X} = [\boldsymbol{X}^1, \ldots, \boldsymbol{X}^B]$ and $\boldsymbol{Y}$ be the predictor matrix and outcome matrix, respectively, that are defined on the same samples $n$. Concretely, the samples are split into $G$ groups $n_1 + n_2 + \cdots + n_G = n$The outcome matrix $\boldsymbol{Y}$ is one-hot-encoded (1/0), with only one column for the binary outcome ($G = 2$) and $G$ columns for the multiclasss outcome ($G \geq 3$).

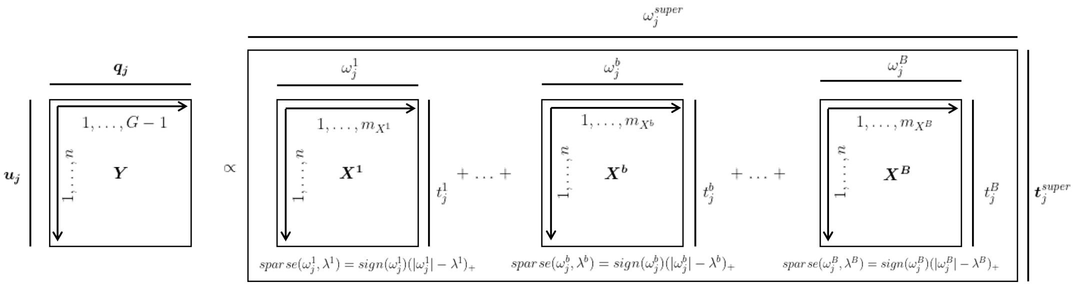


**Fig. S1.** Representation of asmbPLS-DA.

Source: own elaboration.

The objective of asmbPLS-DA is to build a set of orthogonal PLS components such that the covariance between the latent scores $t_j^{super}$ and $u_j$, which represent information from $\boldsymbol{X}$ and $\boldsymbol{Y}$ respectively, is maximized for each PLS component $j = 1, 2, \ldots, J$. Formally, the problem can be stated as an optimization problem.

$$\max_{\{\omega_j^{super}, \omega_j^b, q_j\}} Cov(\boldsymbol{t}_j^{super}, \boldsymbol{u}_j) \tag{S1}$$

$$subject\ to \quad ||\omega_j^b|| = ||\omega_j^{super}|| = 1$$

$$where \quad t_j^b = \frac{\boldsymbol{X}^b \omega_j^b}{\sqrt{m_{X^b}}}, \quad \boldsymbol{T}_j = [t_j^1, \ldots, t_j^B], \quad \boldsymbol{u_j} = \boldsymbol{Y}\boldsymbol{q_j}, \quad \boldsymbol{t}_j^{super} = \boldsymbol{T}_j \omega_j^{super}$$

The optimization algorithm to solve (1) revolves around the following steps for each PLS component $j = 1, \ldots, J$.

- The first dimension reduction is conducted by taking a linear combination of the predictor features (columns) to obtain each of the block scores $t_j^b = \frac{\boldsymbol{X}^b \omega_j^b}{\sqrt{m_{X^b}}}$ for each of the predictor blocks $b = 1, \ldots, B$. Where $\omega_j^b$ contains the block feature weight in $\boldsymbol{X}^b$ which indicates the relevance of the feature to classify between groups, and $m_{X^b}$ is the number of variables in $X^b$ for block scaling.
- Within each block, the soft tresholding function $sparse(\omega_j^b, \lambda^b) = sign(\omega_j^b)(|\omega_j^b| - \lambda^b)$ is implemented as a sparsity criteria which sets as 0 all weights below a quantile treshold $\lambda^b \in [0, 1]$ selected by Cross Validation. By doing this, only the most relevant features will be retained.

- All block scores are combined into $\boldsymbol{T}_j = [t_j^1, \ldots, t_j^B]$ and a new dimension reduction is implemented again by taking a linear combination of the different block scores $\boldsymbol{t}_j^{super} = \boldsymbol{T}_j \omega_j^{super}$, where $\omega_j^{super}$ is the relevance of each block of the matrix $\boldsymbol{X}$ in classifying, while $\boldsymbol{t}_j^{super}$ is the super score that represents the information contained in predictor $\boldsymbol{X}$.
- Finally, the summary vector $\boldsymbol{u}_j$ contains the information on the response $\boldsymbol{Y}$ and $q_j$ is the weight of each column of $\boldsymbol{Y}$, hence $\boldsymbol{u}_j = \boldsymbol{Y} q_j$ is computed.

Once all parameters are estimated $\{\omega_j^{super}, \omega_j^b, q_j\}$ for the first PLS component, $\boldsymbol{X}$ and $\boldsymbol{Y}$ are deflated, and then the deflated matrices are used again for the calculation of the second PLS component, and so on. With that a prediction of $\boldsymbol{Y}$ for a new set of predictors $\boldsymbol{X}'$ are computed through a linear combination.

$$\boldsymbol{Y}' = \boldsymbol{t}_1^{super'} q_1 + \boldsymbol{t}_2^{super'} q_2 + \cdots + \boldsymbol{t}_J^{super'} q_J \tag{S2}$$

With the estimated numeric prediction $\boldsymbol{Y}'$ different decision rules to discriminate between classes, and assign one of the classes $1, 2, \ldots, G$, are implemented in asmbPLS-DA from: fixed cut-off, euclidean distance, mahalanobis distance, principal component analysis and mahalanobis distance, etc.

Note that many solutions can arise from (1) based on the selection of the number of PLS components $J$ and the quantile treshold for each block $\lambda = [\lambda^1, \ldots, \lambda^B]$, to select the optimal value for the hyperparameters a Cross Validation with K-fold is implemented in asmbPLS-DA. The K-fold CV selects the best hyperparameters for which the highest balanced accuracy (BA) is attained. The samples are randomly placed into $K$ groups with the ratios of the samples from different classes being equal in all the groups. Such step is repeated $N_{CV}$ times to generate $N_{CV}$ set of groups. For each of the $K$ groups the last dataset is used for validation while the rest $K-1$ are used for training. The average $K$-fold BA is computed as an average over the $N_{CV}$ sets:

$$BA_{N_{CV},K} = \frac{1}{N_{CV}} \sum_{n_{CV}=1}^{N_{CV}} \frac{1}{K} \sum_{k=1}^{K} BA_{n_{CV},k} \tag{S3}$$

Where $BA_{n_{CV},k}$ is the balanced accuracy using samples from the $k-th$ group of $n_{CV}-th$ set as the validation, the model with the lowest (3) is chosen. The optimal number $J^*$ of PLS components are determined by initially selecting $J^* = 1$, then check whether including one more component decreases the BA by a treshold $0 < \varepsilon < 0.005$, i.e $BA_{J^*+1} + \varepsilon \leq BA_{J^*}$, if true then set $J^*$ to $J^* + 1$ until the treshold criteria is not attained.

#### B. Gene orthology mapping

Gene orthology mapping between humans and disease models were retrieved from the ENSEMBL (Yates et al., 2019) homology between *Homo sapiens* and *Mus musculus*. Only one-to-one orthologs were retrieved, excluding many-to-many and one-to-many relationships. Entrez gene symbols were mapped to HGNC using biomaRt v2.58.2 (Durinck et al., 2009), which were posteriorely used for singIST workflow.

#### C. asmbPLS-DA parameter tuning and model selection

The optimal model is fitted using Leave-One-Out Cross Validation (LOOCV), due to the small sample size of scRNA-seq data, to select the number of PLS components $J^*$, as well as the quantile combination for each block and PLS component $\lambda_j^b = \text{quantile}\{|\omega_j^b|, \lambda^b\}$, $\lambda^b$ is the hyperparameter for quantile tuning $\lambda^b \in [0,1]$, that is used for the sparse criteria $sparse(\omega_j^b, \lambda_j^b) = sign(\omega_j^b)(|\omega_j^b| - \lambda_j^b)_+$. The LOOCV method aims to identify the most effective quantile sparsity combination, for each PLS component $j$, that yield the highest $F_1$ score.

#### D. Superpathway's score $\widehat{y}_{ik}$ decomposition onto cell $\widehat{\gamma}_{ik}^b$ and gene $\widehat{\delta}_{igk}^b$ contributions

For a generic sample $i$ and number of PLS $J$, the asmbPLS-DA predictor is $\widehat{y}_{ik} = \sum_{j=1}^{J} t_{ij}^{super} q_{kj}^T$. Denote $\boldsymbol{T} = [t_{ij}^1, \ldots, t_{ij}^b, \ldots, t_{ij}^B]$ then $\boldsymbol{t}_{ij}^{super} = (\boldsymbol{T} \times_1 \omega^{super}) = \sum_{b=1}^{B} t_{ij}^b \left(\omega_{bj}^{super}\right)^T$, where $(\boldsymbol{T} \times_1 \omega^{super})$ denotes the n-mode product over the mode 1. Hence plugging that in the predictor:

$$\widehat{y}_{ik} = \sum_{j=1}^{J} t_{ij}^{super} q_{kj}^{T} = \sum_{j=1}^{J} \left[ \sum_{b=1}^{B} t_{ij}^{b} \left( \omega_{bj}^{super} \right)^{T} \right] q_{kj}^{T} = \sum_{j=1}^{J} \left[ \sum_{b=1}^{B} t_{ij}^{b} \left( \omega_{bj}^{super} \right)^{T} q_{kj}^{T} \right] \tag{S4}$$

$$= \sum_{b=1}^{B} \left[ \sum_{j=1}^{J} t_{ij}^{b} \left( \omega_{bj}^{super} \right)^{T} q_{kj}^{T} \right] = \sum_{b=1}^{B} \widehat{\gamma}_{ik}^{b} \tag{S5}$$

To obtain the gene contributions note that the gene score $t_{ij}^{b} = \frac{\mathbf{C}^{b}\omega^{b}}{\sqrt{|\mathcal{G}_{p}^{b}|}} = \sum_{g \in \mathcal{G}_{p}^{b}} \frac{x_{ig}^{b}\omega_{gj}^{b}}{\sqrt{|\mathcal{G}_{p}^{b}|}}$, and plugging such into Eq. (S5):

$$\widehat{y}_{ik} = \sum_{b=1}^{B} \left[ \sum_{j=1}^{J} t_{ij}^{b} \left( \omega_{bj}^{super} \right)^{T} q_{kj}^{T} \right] = \sum_{b=1}^{B} \left[ \sum_{j=1}^{J} \left( \sum_{g \in \mathcal{G}_{p}^{b}} \frac{x_{ig}^{b}\omega_{gj}^{b}}{\sqrt{|\mathcal{G}_{p}^{b}|}} \right) \left( \omega_{bj}^{super} \right)^{T} q_{kj}^{T} \right] \tag{S6}$$

$$= \sum_{b=1}^{B} \left[ \sum_{j=1}^{J} \left( \sum_{g \in \mathcal{G}_{p}^{b}} \frac{x_{ig}^{b}\omega_{gj}^{b}}{\sqrt{|\mathcal{G}_{p}^{b}|}} \left( \omega_{bj}^{super} \right)^{T} q_{kj}^{T} \right) \right] = \sum_{b=1}^{B} \sum_{g \in \mathcal{G}_{p}^{b}} \left[ \sum_{j=1}^{J} \frac{x_{ig}^{b}\omega_{gj}^{b}}{\sqrt{|\mathcal{G}_{p}^{b}|}} \left( \omega_{bj}^{super} \right)^{T} q_{kj}^{T} \right] \tag{S7}$$

Which gets us to $\widehat{y}_{ik} = \sum_{b=1}^{B} \sum_{g \in \mathcal{G}_{p}^{b}} \widehat{\delta}_{igk}^{b}$. To compute predictions the blocks are loading deflated.

#### E. An intuitive explanation of recapitulations

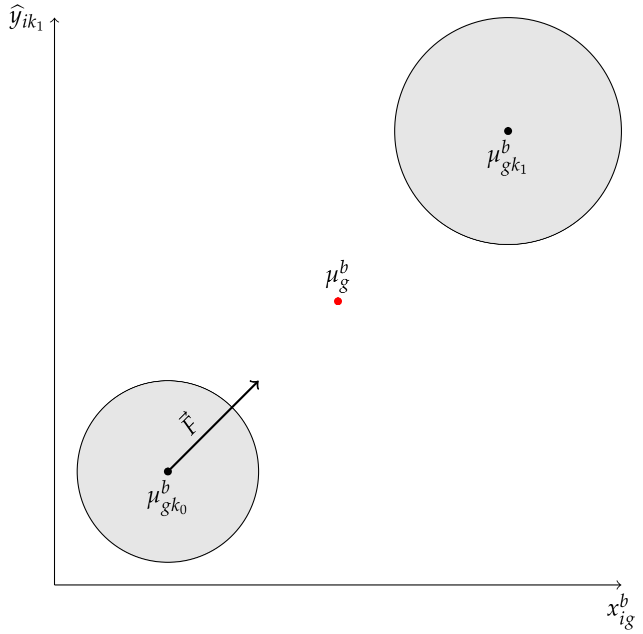


**Fig. S2.** Illustration of singIST recapitulations. Y-axis represents superpathway's score while X-axis is gene expression of a sample for gene $g$ within cell type $b$, circles represent class domain. $\mu_{gk_0}^{b}$ and $\mu_{gk_1}^{b}$ are centroids within class and $\mu_{g}^{b}$ is the pooled gene centroid. The direction and magnitude of movement of base class, for the disease model, is noted as $\vec{F}$ of base class

The direction and magnitude of the change depends on a variety of factors $\vec{F} = f_{\omega^{super},\omega^{b}}\left( \mu_{g}^{b} r_{\widetilde{g}}^{b}, \exists \widetilde{g} \equiv g, \exists b \right)$. On one hand, if the gene does not have an ortholog ($\nexists \widetilde{g} \equiv g$) or cell type does not exist ($\nexists b$) then there is no direction nor magnitude of movement $\vec{F} = 0$, note that this varies from disease model to disease model. Likewise, if either gene or cell type are not relevant at all $CIP^{b} = 0$ or $GIP_{g}^{b} = 0$, then the direction and magnitude of movement will $\vec{F} = 0$, note that this will be true for any disease model. Lastly, if noone of the former cases hold, the direction and magnitude of movement will be defined by $\vec{F} = \mu_g^b r_{\tilde{g}}^b$, the direction comes from the disease model $r_{\tilde{g}}^b$ while the magnitude comes from both $r_{\tilde{g}}^b$ and $\mu_g^b$, the quantity $\mu_g^b r_{\tilde{g}}^b$ gives us the expected change in the human gene expression space if those were to behave like the differences observed in the disease model, measured through $r_{\tilde{g}}^b$. Once $\vec{F}$ is applied, the closer the base samples classes are to the target class samples, the higher the recapitulation.

##### F. Cell type recapitulation as sum of gene contributions $\widehat{\Gamma f}_{k_1}^b = \sum_g \widehat{\Delta f}_{gk_1}^b$

Since $\ell(x_{ig}^b, r_{\tilde{g}}^b; \mu_g^b)$ defines an affine transformation, the change $x_{ig}^b{}' - x_{ig}^b$ is constant for all original samples $i \in K_0 := \{1 \leq i \leq n | y_{ik_0} = 1\}$, further the predictor $\widehat{y}_{ik}$ is a linear transformation, thus the change $\widehat{y}_{ik}' - \widehat{y}_{ik}$ is also constant for all original samples $i \in K_0$. As shown in Eq. (S5) the predictor $\widehat{y}_{ik}$ can be decomposed onto cell contributions $\widehat{y}_{ik}' - \widehat{y}_{ik} = \sum_{b=1}^B \left( \widehat{\gamma}_{ik}^b{}' - \widehat{\gamma}_{ik}^b \right)$, for which there $\exists \{\widetilde{\gamma}_k^b\}_{b,k}$ constants such that $\widehat{y}_{ik}' - \widehat{y}_{ik} = \sum_{b=1}^B \widetilde{\gamma}_k^b$. Without loss of generality fix a $b \in \{1, \ldots, B\}$, the constant difference in cell contribution is $\widehat{\gamma}_{ik}^b{}' = \widetilde{\gamma}_k^b + \widehat{\gamma}_{ik}^b$, note that applying $\underset{i \in K_0}{\text{median}} \left( \widehat{\gamma}_{ik}^b{}' \right) = \underset{i \in K_0}{\text{median}} \left( \widetilde{\gamma}_k^b + \widehat{\gamma}_{ik}^b \right)$ since *singIST treated samples* do not belong to $K_0$ samples $\widehat{\gamma}_{ik}^b{}' = \widetilde{\gamma}_k^b + \underset{i \in K_0}{\text{median}} \left( \widehat{\gamma}_{ik}^b \right)$, plugging it into the observed cell type recapitulation:

$$\widehat{\Gamma}_{k_1}^b = \underset{i \in K_1}{\text{median}} \left( \widehat{\gamma}_{ik}^b{}' \right) - \underset{i \in K_0}{\text{median}} \left( \widehat{\gamma}_{ik}^b \right) = \underset{i \in K_1}{\text{median}} \left( \widetilde{\gamma}_{k_1}^b + \underset{i \in K_0}{\text{median}} \left( \widehat{\gamma}_{ik}^b \right) \right) - \underset{i \in K_0}{\text{median}} \left( \widehat{\gamma}_{ik}^b \right) \quad \text{(S8)}$$

$$= \widetilde{\gamma}_{k_1}^b \quad \text{(S9)}$$

With the same reasoning, there also $\exists \{\widehat{\Delta}_{gk}^b\}_{g,k,b}$ such that $\widetilde{\gamma}_k^b = \sum_{g \in \mathcal{G}_p^b} \widehat{\Delta}_{gk}^b$ , which is straightforward then that $\widehat{\Gamma f}_{k_1}^b = \sum_{g \in \mathcal{G}_p^b} \widehat{\Delta f}_{gk_1}^b$.

##### G. Recapitulations as a function of $\omega^{super}$, $\omega^b$, $\exists b$, $\exists \tilde{g} \equiv g$, $r_{\tilde{g}}^b$

For a fixed $b$ and $g$, we start by explicitly computing the absolute gene contribution to cell type recapitulation $\widehat{\Delta}_{gk_1}^b := \widehat{\delta}_{igk_1}^b{}' - \widehat{\delta}_{igk_0}^b$:

$$\widehat{\Delta}_{gk_1}^b = \sum_{j=1}^J \frac{q_{k_1 j}^T}{\sqrt{|\mathcal{G}_p^b|}} \left( \omega_{bj}^{super} \right)^T \omega_{gj}^b \left( x_{ig}^b{}' - x_{ig}^b \right) \quad \text{(S10)}$$

If $\nexists b$ and/or $\nexists \tilde{g} \equiv g$ by construction $x_{ig}^b{}' = x_{ig}^b$, hence $\widehat{\Delta}_{gk_1}^b = 0$ which yields to $\widehat{\Delta f}_{gk_1}^b = 0$. Similarly, assume $GIP_g^b = 0$ which implies that $\omega_{gj}^b = 0 \quad \forall j$, then $\widehat{\Delta}_{gk_1}^b = 0$ indistinguisably of $x_{ig}^b{}' - x_{ig}^b$. Likewise, if a cell type is not important in human condition $CIP^b = 0$ it implies that $\omega_{bj}^{super} = 0 \quad \forall j$ , hence $\widehat{\Gamma}_{k_1}^b = \sum_{g \in \mathcal{G}_p^b} \widehat{\Delta}_{gk_1}^b = 0$ and the cell type recapitulation becomes null $\widehat{\Gamma f}_{k_1}^b = 0$.

Assume a non trivial case where $CIP^b, GIP_g^b > 0$ and conditions $\exists b, \exists \tilde{g} \equiv g, C = \min_{i \in \{1 \leq i \leq n | y_{ik_1} = 1\}} \{x_{ig}^b + \mu_g^b r_{\tilde{g}}^b\} \geq 0$ hold true, then Eq. (S11) becomes:

$$\widehat{\Delta}_{gk_1}^b = \frac{\mu_g^b}{\sqrt{|\mathcal{G}_p^b|}} r_{\tilde{g}}^b \sum_{j=1}^J q_{k_1 j}^T \left( \omega_{bj}^{super} \right)^T \omega_{gj}^b \quad \text{(S11)}$$

Note that, $\mu_g^b$, $|\mathcal{G}_p^b|$ are always positive and so are $q_{k_1 j}^T$ and $\left( \omega_{bj}^{super} \right)^T$ when class is one-hot-encoded, hence the direction of $\widehat{\Delta}_{gk_1}^b$ as $sign \left( \widehat{\Delta}_{gk_1}^b \right) = sign \left( \sum_{j=1}^J q_{k_1 j}^T \left( \omega_{bj}^{super} \right)^T \omega_{gj}^b \right) sign \left( r_{\tilde{g}}^b \right)$, as the agreement in direction between the disease model estimated FC and the estimated direction of the gene within the cell type for humans. Note that differences between disease models' recapitulation will be based on their orthology, cell type agreement and estimated FC, as all other parameters are fixed by disease model.

In a similar fashion, it can be proven that pathway recapitulation is constant and can be derived as the sum of cell type recapitulations, which eventually yields to the same conclusions.

##### H. Parameters variabilities and significance of the optimal asmbPLS-DA

First, we prove that $\sum_{g \in \mathcal{G}_p^b} GIP_g^b = 1$. From the optimization problem of asmbPLS-DA it stems that $||\omega_{\cdot j}^b|| = 1$.

$$\sum_{g \in \mathcal{G}_p^b} GIP_g^b = \frac{\sum_{g \in \mathcal{G}_p^b} \sum_{j=1}^{J^*} q_{k_1 j} \left(\omega_{gj}^b\right)^2}{\sum_{j=1}^{J^*} q_{k_1 j}} = \frac{\sum_{j=1}^{J^*} \left[ q_{k_1 j} \sum_{g \in \mathcal{G}_p^b} \left(\omega_{gj}^b\right)^2 \right]}{\sum_{j=1}^{J^*} q_{k_1 j}} = \frac{\sum_{j=1}^{J^*} q_{k_1 j} ||\omega_{\cdot j}^b||^2}{\sum_{j=1}^{J^*} q_{k_1 j}} = 1$$

We now detail the rationale of Bonferroni's multiple testing correction of $GIP_g^b$ significance test by using $m_0^b = \left\lfloor \prod_{j=1}^{J^*} \lambda_j^b |\mathcal{G}_p^b| \right\rfloor$ as the minimum number of true hypothesis. For cell type $b$, let $H_{0,1}^b, \ldots, H_{0,g}^b, \ldots, H_{0,g_b}^b$ be the family of null hypothesis defined as $H_{0,g}^b$ : median$(\mathcal{D}_{g,Jackknife}^b) \leq$ median$(\mathcal{D}_{0,g}^b)$ and their corresponding p-values $p_1^b, \ldots, p_g^b, \ldots, p_{g_b}^b$. $|\mathcal{G}_p^b|$ are the total number of hypothesis to be tested and $M_0^b$ the number of true null hypothesis. We provide a lower bound of $M_0^b$ by noting that for any cell with quantile sparsity values $\{\lambda_j^b\}_{j=1}^{J^*}$ it holds that $P_{\mathcal{D}}(GIP_g^b = 0) = \prod_{j=1}^{J^*} \lambda_j^b$ for any distribution $\mathcal{D}$, and non-trivial gene $x_{ig}^b = 0, \forall i$. asmbPLS-DA retains the top $\lambda_j^b \in [0,1]$ weights, hence by definition of the sparsity criteria and orthogonality of PLS components $P_{\mathcal{D}}(\{\omega_{gj}^b = 0 : \forall j\}) = \prod_{j=1}^{J^*} \lambda_j^b$ for any non-trivial gene $g$. Without loss of generality assume the weights $q_{k_1 j} \geq 0$, which is always the case when the class is one-hot-encoded and/or response scale is greater than 0.

$$P_{\mathcal{D}}\left(GIP_g^b = 0 | q_{k_1 j} \geq 0, \forall j\right) = P_{\mathcal{D}}(\{\omega_{gj}^b = 0 : \forall j\}) = \prod_{j=1}^{J^*} \lambda_j^b, \quad \forall \mathcal{D}$$

In conclusion, for at least $\left\lfloor \prod_{j=1}^{J^*} \lambda_j^b |\mathcal{G}_p^b| \right\rfloor$ tests the null hypothesis will be true due to the sparsity criteria. The lower bound of the number of true null hypothesis $M_0^b \geq m_0^b = \left\lfloor \prod_{j=1}^{J^*} \lambda_j^b |\mathcal{G}_p^b| \right\rfloor$ serves as a correction value to preserve a confidence treshold of $\alpha$, if $\left\lfloor \prod_{j=1}^{J^*} \lambda_j^b \right\rfloor = 0$ then $m_0^b$ was set to $m_0^b = 1$

$$FWER_{Bonferroni} \leq \frac{m_0^b}{|\mathcal{G}_p^b|} \alpha \leq \alpha$$

Which trivially holds true due to the chain of inequalities:

$$\alpha \geq \frac{M_0^b}{|\mathcal{G}_p^b|} \alpha \geq \frac{m_0^b}{|\mathcal{G}_p^b|} \alpha \geq \frac{\prod_{j=1}^{J^*} \lambda_j^b |\mathcal{G}_p^b|}{|\mathcal{G}_p^b|} \alpha = \prod_{j=1}^{J^*} \lambda_j^b \alpha$$
