## Supplementary Material S2 for "singIST: an integrative method for comparative single-cell transcriptomics between disease models and humans"

| Organism | Cluster | Cell type granularity 1** | Cell type granularity 2*** | Action | Reason |
|---|---|---|---|---|---|
| *Homo Sapiens Sapiens** | KC-1 | Keratinocytes | Suprabasal Keratinocyte | KEEP GRANULARITY 1 | No relevant information reported on granularity 2 Keratinocytes in the original paper, only for granularity 1 Keratinocyte |
| | KC-2 | | Basal Keratinocyte | | |
| | KC-3 | | Late differentiation Keratinocyte | | |
| | KC-4 | | Proliferating Keratinocyte | | |
| | KC-5 | | ? | | |
| | Tregs | T-cell | T-regs | KEEP GRANULARITY 2 | |
| | TC-1 | T-cell | Tissue Resident Memory T-cell | | |
| | TC-2 | T-cell | CD8+ effector T-cell | | |
| | TC-3 | T-cell | (CD161)+ T-cell / Th2a ? | DROP CLUSTER | - Small cluster according to [1] "*The smaller clusters TC-3, TC-4, and TC-5 were either absent (TC-3) or only detectable in small numbers (TC-4 and TC-5) in healthy control samples* "<br>- Not clear from the publication if its (CD161)+ T-cell or Th2A |
| | TC-4 | T-cell | ? | DROP CLUSTER | Non-identified cell type |
| | TC-5 | T-cell | Proliferating T-cell | DROP CLUSTER | Small cluster according to [1] "*The smaller clusters TC-3, TC-4, and TC-5 were either absent (TC-3) or only detectable in small numbers (TC-4 and TC-5) in healthy control samples* " |
| | TC-6 | T-cell | Natural Killer T-cell | KEEP GRANULARITY 2 | |
| | Melanocytes | Melanocytes | | KEEP GRANULARITY 1 | It's the only granularity reported |
| | LC | Dendritic cells | Langerhans cells | KEEP GRANULARITY 2 | |
| | DC-1 | Dendritic cells | Myeloid cells | | |
| | DC-2 | Dendritic cells | Mature DCs | DROP CLUSTER | - Not relevant according to [1] "*Although the list of differentially expressed genes in DC-2 was relatively short, likely due to very small cell numbers and thus lacking statistical power, a few anti-inflammatory genes showed marked up-regulation during treatment, especially,...* "<br>- We are not considering treatment period, only HC and Baseline.<br>- Very small cluster to consider it for analysis |
| | DC-3 | Dendritic cells | Plasmacytoid DCs | DROP CLUSTER | - Very small population according to [1] "*We found a very small population of plasmacytoid DCs (DC-3)...* " |
| | MastC_Others | ? | | DROP CLUSTER | Non-identified cell type |

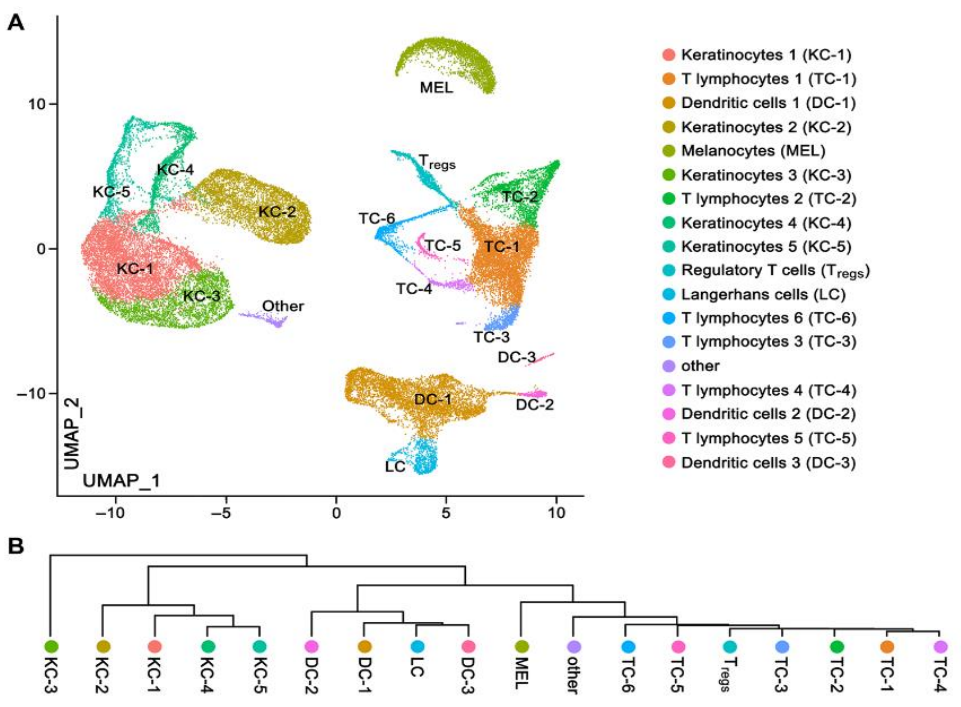


*Human data extracted from [1] "*Persistence of mature dendritic cells, TH2A, and Tc2 cells characterize clinically resolved atopic dermatitis under IL-4Ralpha blockade* " url: https://www.science.org/doi/epdf/10.1126/sciimmunol.abe2749

** If "?" cell type granularity 1 is not clearly identified in the publication

*** If "?" cell type granularity 2 is not clearly identified in the publication

| Organism | Disease model(s) | Cluster | Cell type granularity 1* | Cell type granularity 2* | Action | Reason |
|---|---|---|---|---|---|---|
| *Mus musculus* | *Imiquimod (5%)* and *Oxazolona 3 Challenges* | 0 | Mac (Macrophages) | | | |
| | | 1 | DETC (Dendritic Epidermal T-cells) | | | |
| | | 2 | M/MdM (Monocyte derived Macrophages) | | | |
| | | 3 | dgammadeltaT | | | |
| | | 4 | cDC2 (Type 2 conventional Dendritic cell) | | | |
| | | 5 | ILC2 (Type 2 Innate Lymphoid cell) | | | |
| | | 6 | Thet (T-cell heterogeneous) | Treg<br>Tconv (Conventional T-cell)<br>CD8+ (CD8+ effector T-cell)<br>DNT (Double Negative T-cell) | | |
| | | 7 | Fibroblasts1 | | | |
| | | 8 | Indeterminate1 | | | |
| | | 9 | M/B (Mast cells and Basophils) | | | |
| | | 10 | Neu (Neutrophils) | | | |
| | | 11 | Cell cycle1 | | | |
| | | 12 | Keratinocytes | | | |
| | | 13 | Cell cycle2 | | | |
| | | 14 | cDC1 (Type 1 conventional Dendritic cell) | | | |
| | | 15 | LC (Langerhans cell) | | | |
| | | 16 | mDCs (Mature Dendritic cell) | | | |
| | | 17 | NK (Natural Killer cell) | | | |
| | | 18 | Endothelial | | | |
| | | 19 | Indeterminate2 | | | |
| | | 20 | Fibroblasts2 | | | |

*IMQ-OXA data extracted from [2] "Single-Cell Profiling Reveals Divergent, Globally Patterned Immune Responses in Murine Skin Inflammation" url: https://pubmed.ncbi.nlm.nih.gov/33205009/

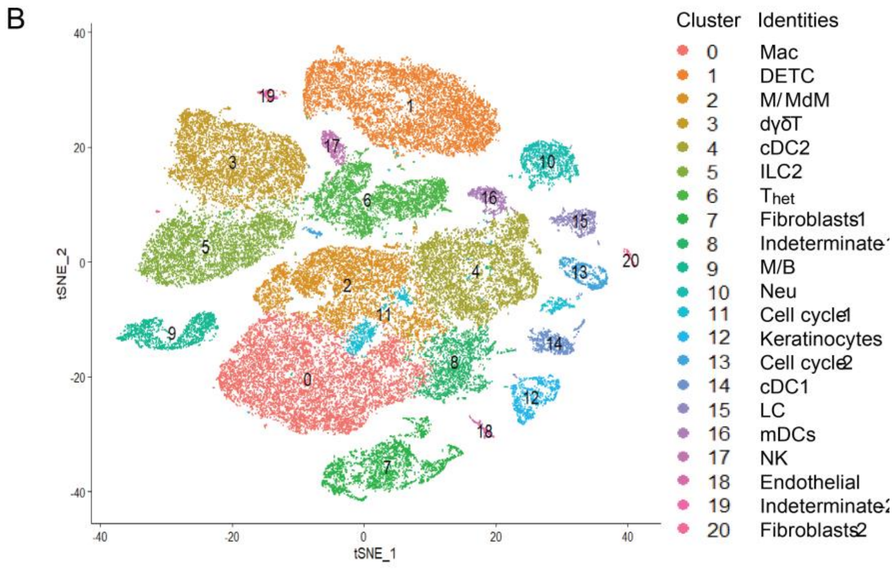


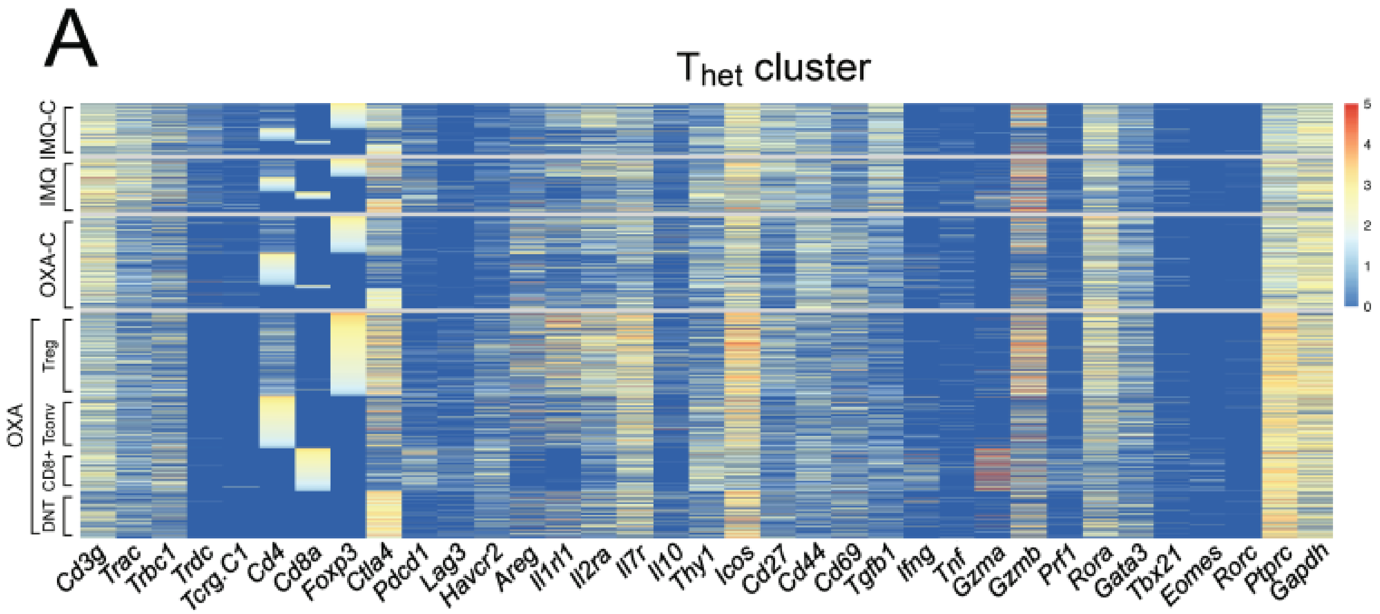


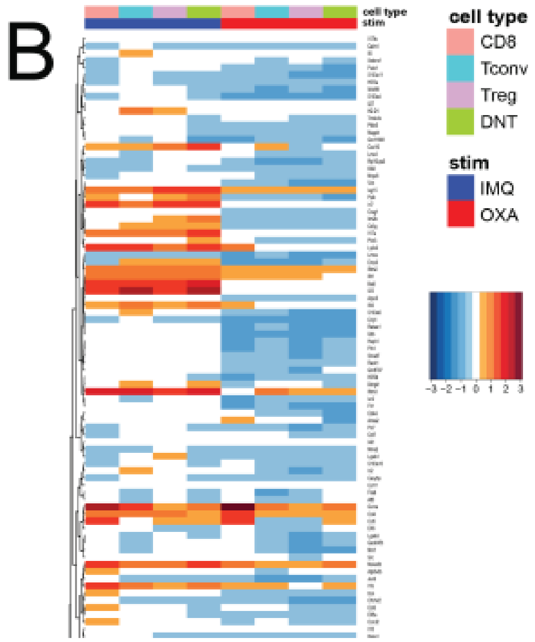

| **Human cells** | **IMQ-OXA cluster** | **IMQ-OXA cells** | **Reason** |
|---|---|---|---|
| Keratinocytes | 12 | Keratinocytes | |
| Tregs | 6 | T-heterogeneous (Treg) | |
| Tissue Resident Memory T-cell | **?** | **?** | |
| CD8+ effector T-cell | 6 | T-heterogeneous (CD8+ effector T-cell) | |
| Natural Killer T-cell | 17 | NK | |
| Melanocytes | | **None** | Melanocytes are not present in murine ear skin but either bulge and bulb regions of hair follicles, tail or ventral paws of non-hairy skin according to [3] "*Firstly, mouse pelage skin interfollicular epidermis entirely lacks functional, pigment-producing melanocytes. While murine melanocytes are found either in the bulge and bulb regions of hair follicles, in the tail, or in the ventral paws of non-hairy mouse skin, functional human melanocytes are mostly located in the basal layer of the epidermis (Gola et al., 2012).* " |
| Langerhans cells | 15 | LC (Langerhans cells) | |
| Myeloid cells | 0, 2, 9, 10, 17 | Mac (Macrophages), M/MdM (Monocyte derived Macrophages), M/B (Mast cells/Basophils), Neu (Neutrophils) | |

[3] "*Characterization of a melanocyte progenitor population in human interfollicular epidermis* ". Cell Reports.
