## Supplementary Material S3 for "singIST: an integrative method for comparative single-cell transcriptomics between disease models and humans"

### Supplementary Material S3: Results

**Table S1.** Characteristics of fitted optimal asmbPLS-DA for all superpathways.

| Pathway | $J^{*a}$ | Gene set size[b] | adj pvalue[c] | Dendritic Cell[d] | | Keratinocyte | | Langerhans Cell | | Melanocyte | | T-cell | |
|---|---|---|---|---|---|---|---|---|---|---|---|---|---|
| | | | | $\lambda^1$ | $GIP_g^1*$ | $\lambda^2$ | $GIP_g^2*$ | $\lambda^3$ | $GIP_g^3*$ | $\lambda^4$ | $GIP_g^4*$ | $\lambda^5$ | $GIP_g^5*$ |
| Cytokine-Cytokine receptor interaction [KEGG] | 2 | 263 | $p \leq 0.001$ | 0.75<br>0.05 | 31 | 0.95<br>0.05 | 18 | 0.55<br>0.05 | 50 | 0.95<br>0.65 | 28 | 0.95<br>0.45 | 17 |
| Chemokine receptors bind chemokines [REACTOME] | 1 | 57 | $p \leq 0.001$ | 0.05 | 13 | 0.05 | 20 | 0.45 | 9 | 0.95 | 3 | 0.85 | 7 |
| Chemokine signaling pathway [KEGG] | 1 | 187 | $p = 0.065$ | 0.05 | 35 | 0.05 | 44 | 0.05 | 37 | 0.05 | 41 | 0.05 | 39 |
| Inflammation pathway [BIOCARTA] | 1 | 27 | $p \leq 0.05$ | 0.05 | 9 | 0.05 | 8 | 0.75 | 4 | 0.45 | 5 | 0.05 | 8 |
| Th1/Th2 Differentiation [BIOCARTA] | 1 | 21 | $p = 0.065$ | 0.05 | 5 | 0.05 | 8 | 0.05 | 7 | 0.05 | 7 | 0.05 | 6 |
| Cytokine Network [BIOCARTA] | 1 | 19 | $p \leq 0.05$ | 0.05 | 6 | 0.05 | 5 | 0.05 | 6 | 0.05 | 8 | 0.05 | 5 |
| Dendritic Cells in Th1/Th2 Development [BIOCARTA] | 1 | 17 | $p \leq 0.05$ | 0.55 | 3 | 0.95 | 1 | 0.95 | 1 | 0.45 | 4 | 0.35 | 4 |
| JAK-STAT signaling pathway [KEGG] | 1 | 155 | $p = 0.065$ | 0.05 | 34 | 0.05 | 38 | 0.05 | 32 | 0.05 | 41 | 0.05 | 32 |
| Asthma [KEGG] | 1 | 28 | $p = 0.065$ | 0.05 | 8 | 0.05 | 11 | 0.95 | 2 | 0.95 | 2 | 0.05 | 8 |
| Toll-like receptor signaling pathway [KEGG] | 1 | 102 | $p = 0.067$ | 0.05 | 21 | 0.05 | 30 | 0.05 | 23 | 0.05 | 24 | 0.05 | 22 |
| IL12 signaling mediated by STAT4 [PID] | 2 | 32 | $p = 0.052$ | 0.05<br>0.85 | 6 | 0.05<br>0.85 | 7 | 0.05<br>0.95 | 5 | 0.05<br>0.95 | 7 | 0.05<br>0.45 | 10 |
| CD40/CD40L signaling [PID] | 1 | 15 | $p = 0.067$ | 0.05 | 3 | 0.05 | 5 | 0.05 | 5 | 0.05 | 4 | 0.05 | 5 |
| IL4-mediated signaling events [PID] | 2 | 64 | $p = 0.052$ | 0.05<br>0.45 | 19 | 0.05<br>0.85 | 13 | 0.05<br>0.85 | 14 | 0.05<br>0.95 | 16 | 0.05<br>0.95 | 11 |
| IL23-mediated signaling events [PID] | 2 | 37 | $p = 0.052$ | 0.05<br>0.05 | 10 | 0.05<br>0.95 | 8 | 0.05<br>0.25 | 13 | 0.05<br>0.85 | 9 | 0.05<br>0.05 | 10 |
| CXCR3-mediated signaling events [PID] | 1 | 43 | $p = 0.053$ | 0.05 | 9 | 0.25 | 12 | 0.05 | 8 | 0.95 | 3 | 0.55 | 9 |
| Hematopoietic cell lineage [KEGG] | 2 | 85 | $p = 0.052$ | 0.05<br>0.25 | 23 | 0.05<br>0.95 | 21 | 0.35<br>0.95 | 15 | 0.05<br>0.95 | 15 | 0.35<br>0.05 | 26 |
| IL2 signaling events mediated by STAT5 [PID] | 1 | 30 | $p = 0.052$ | 0.75 | 3 | 0.95 | 2 | 0.65 | 7 | 0.75 | 4 | 0.85 | 4 |
| NOD-like receptor signaling pathway [KEGG] | 1 | 62 | $p \leq 0.05$ | 0.75 | 10 | 0.85 | 8 | 0.95 | 3 | 0.75 | 10 | 0.95 | 4 |
| Downstream signaling in naïve CD8+ T cells [PID] | 2 | 65 | $p \leq 0.001$ | 0.05<br>0.55 | 20 | 0.05<br>0.95 | 18 | 0.05<br>0.95 | 18 | 0.45<br>0.85 | 11 | 0.95<br>0.05 | 20 |
| T cell receptor signaling pathway [KEGG] | 1 | 108 | $p = 0.052$ | 0.05 | 20 | 0.05 | 27 | 0.05 | 23 | 0.15 | 20 | 0.25 | 20 |
| Cytokine signaling in Immune system [REACTOME] | 2 | 701 | $p = 0.052$ | 0.05<br>0.05 | 199 | 0.05<br>0.05 | 232 | 0.05<br>0.85 | 119 | 0.05<br>0.05 | 221 | 0.05<br>0.05 | 200 |
| Signaling by Interleukins [REACTOME] | 1 | 449 | $p = 0.052$ | 0.95 | 20 | 0.85 | 23 | 0.65 | 91 | 0.95 | 19 | 0.85 | 52 |

$a$. Optimal number of PLS components, for LOOCV was set to $J \leq 3$, $b$. Gene set size $c$. Adj. p-val of global significance, $d$. $\lambda^b$ optimal quantiles of cell type $b$. The quantile space were set to 100000 combinations of $\lambda^b$ values ranging along {0.05, …, 0.55, …, 0.95}. $GIP_g^b*$ number of statistically significant $GIP_g^b$ whose $adj.p - value \leq 0.05$, permutation tests were run on 10000 permutations.

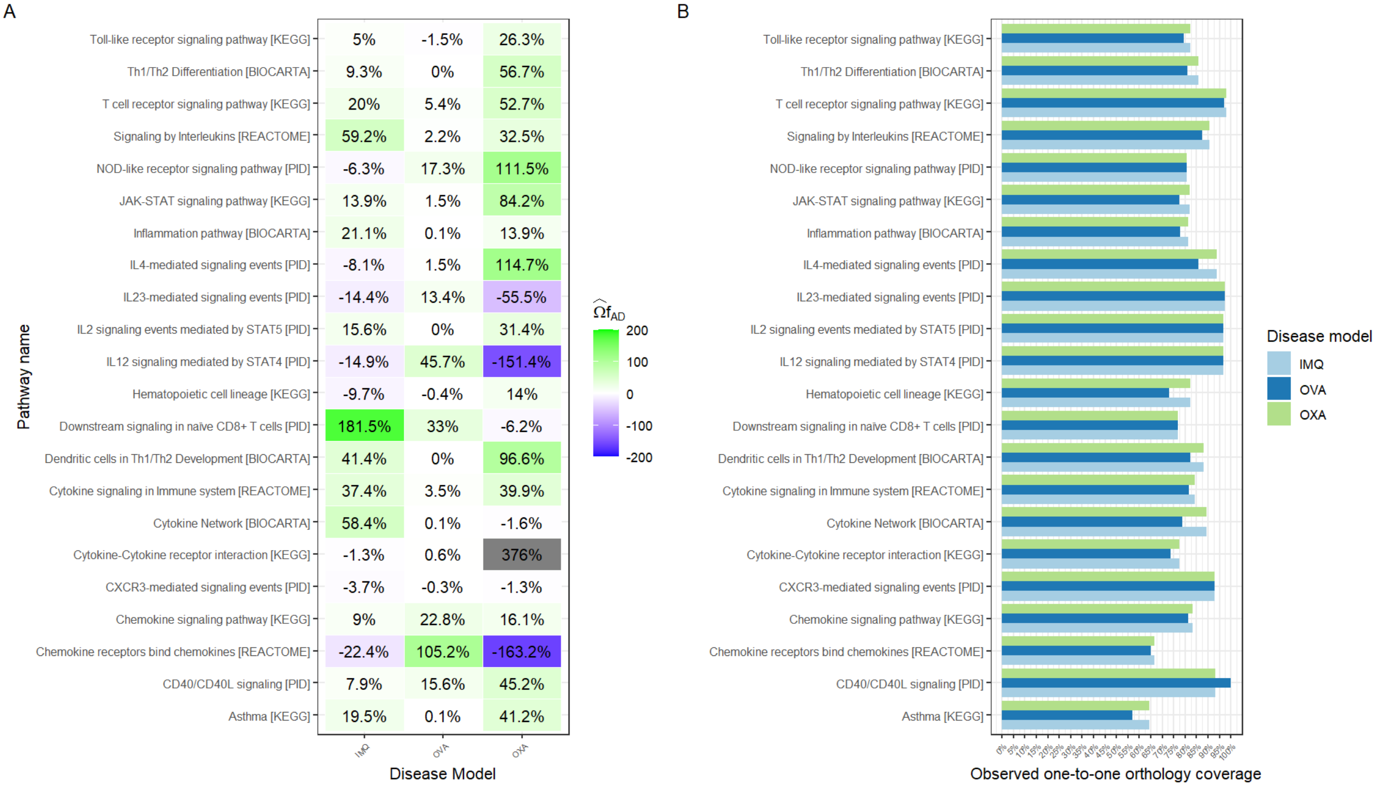


**Fig. S1. Superpathway recapitulation and observed one-to-one orthology of AD disease models. A)** Superpathway predicted recapitulation as a fraction of the superpathway reference recapitulation for IMQ, OXA and OVA across all pathways under study. Negative recapitulations refer to opposed directions with human observed condition, while positive recapitulations define agreement in direction. **B)** Observed one-to-one orthology coverage refers to number of observed and one-to-one ortholog genes in disease model as a fraction of pathway gene set size. Despite all disease models belong to the same organism *mus musculus* their differences in observed orthology one-to-one coverage come from sequenced reads.

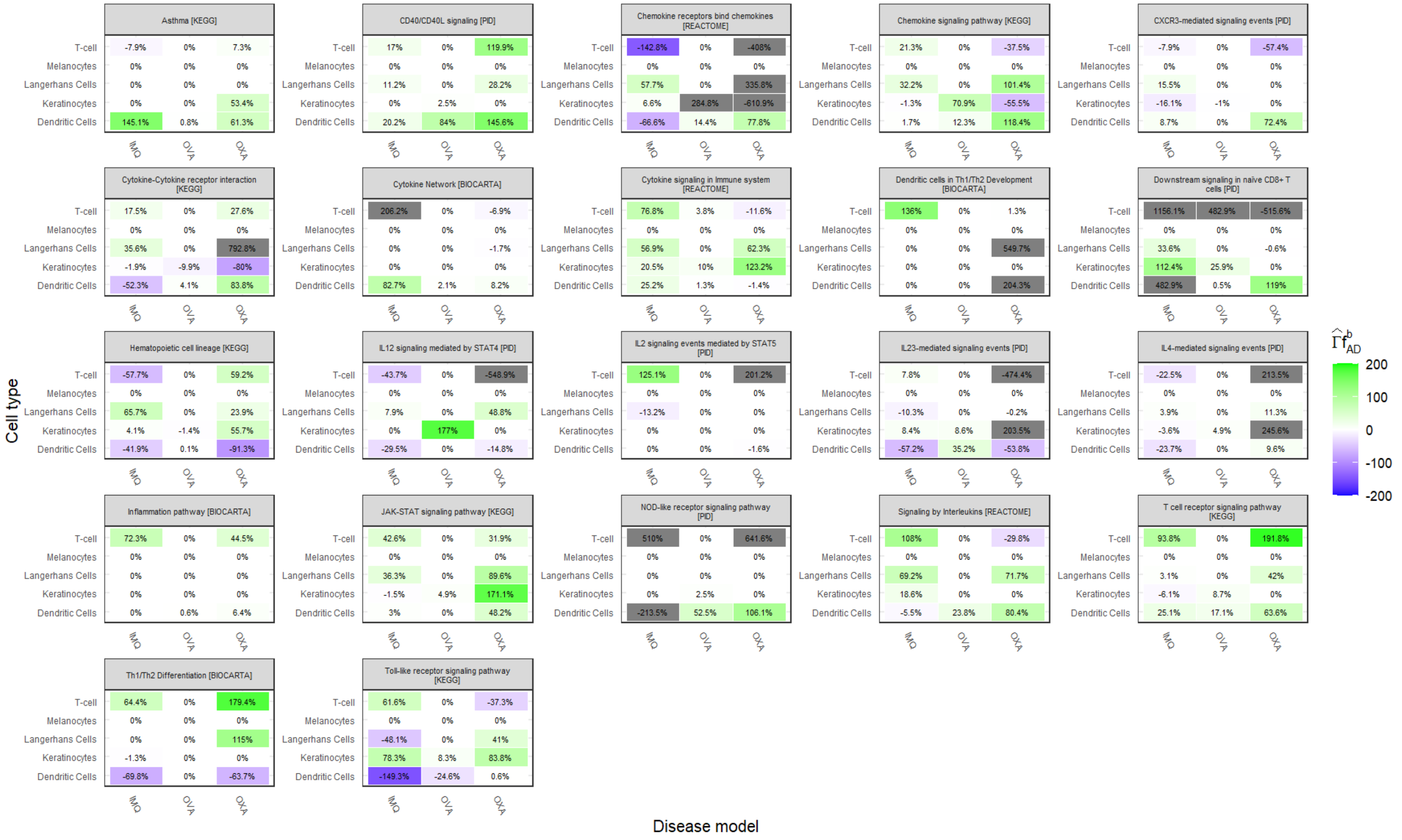


**Fig. S2.** Cell type recapitulation for all AD disease models and pathways under analysis.

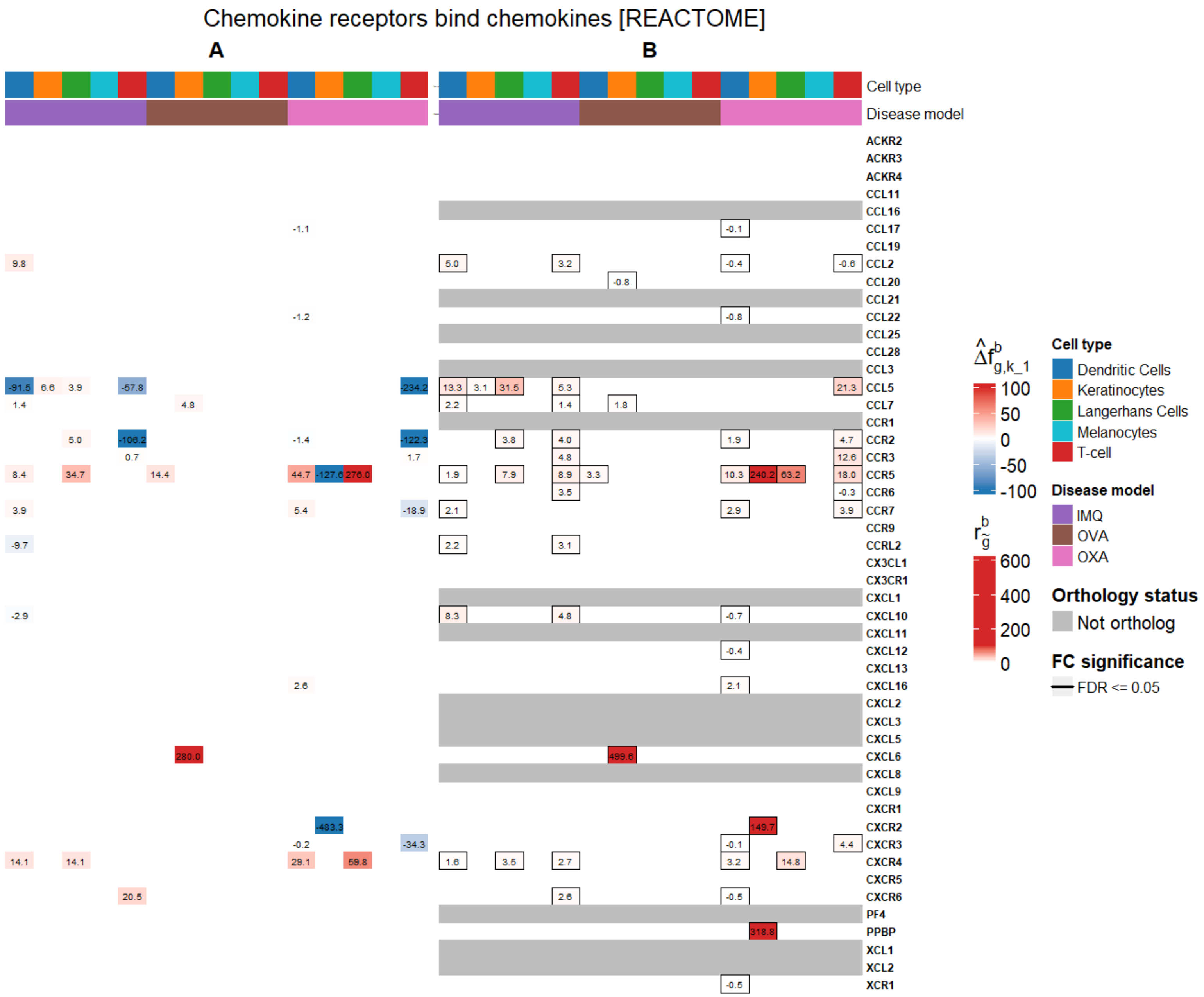
Chemokine receptors bind chemokines [REACTOME]
A
B
Cell type
Disease model
ACKR2
ACKR3
ACKR4
CCL11
CCL16
CCL17
CCL19
CCL2
CCL20
CCL21
CCL22
CCL25
CCL28
CCL3
CCL5
CCL7
CCR1
CCR2
CCR3
CCR5
CCR6
CCR7
CCR9
CCRL2
CX3CL1
CX3CR1
CXCL1
CXCL10
CXCL11
CXCL12
CXCL13
CXCL16
CXCL2
CXCL3
CXCL5
CXCL6
CXCL8
CXCL9
CXCR1
CXCR2
CXCR3
CXCR4
CXCR5
CXCR6
PF4
PPBP
XCL1
XCL2
XCR1
Cell type
Dendritic Cells
Keratinocytes
Langerhans Cells
Melanocytes
T-cell
Disease model
IMQ
OVA
OXA
Orthology status
Not ortholog
FC significance
FDR <= 0.05

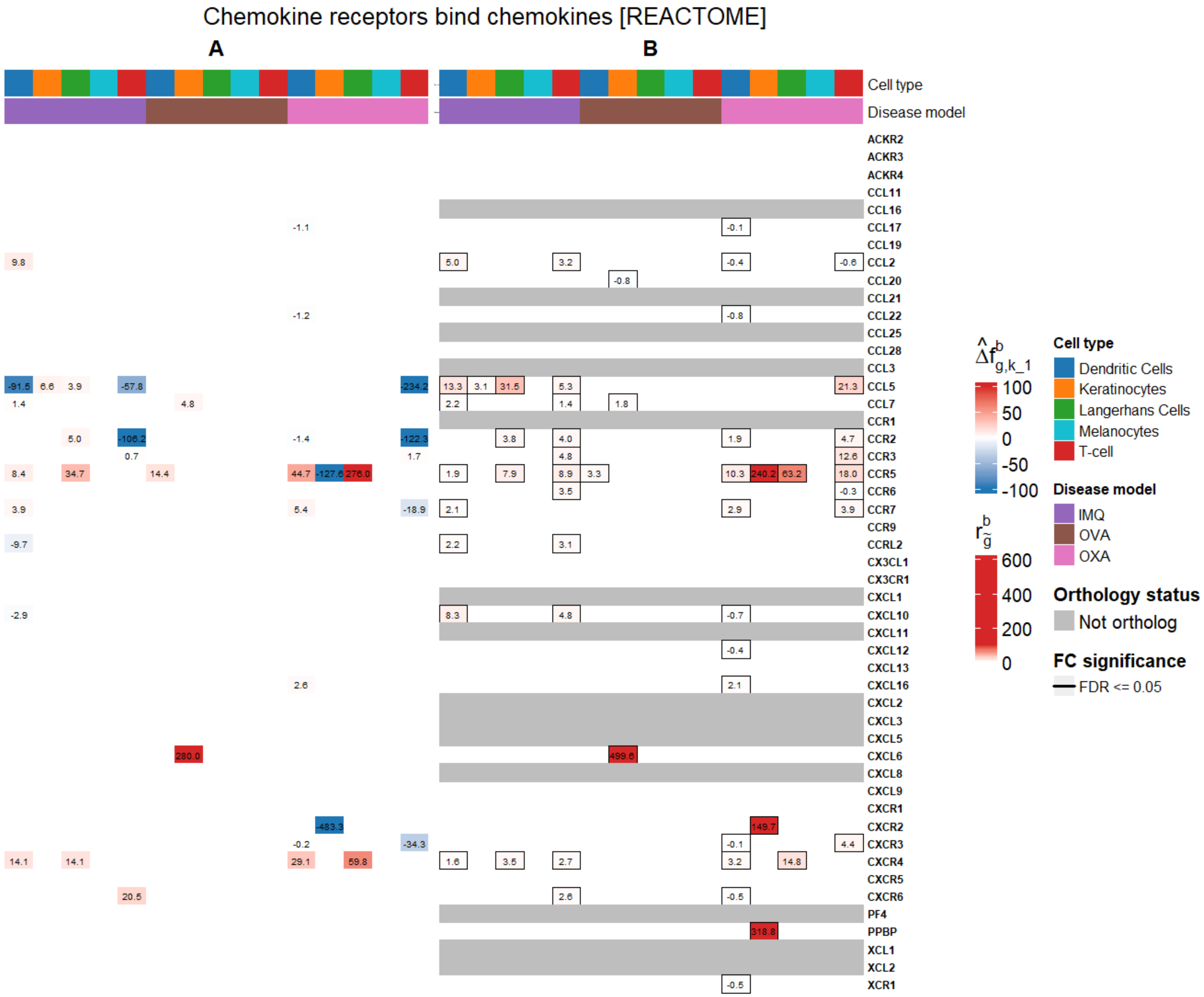

Chemokine receptors bind chemokines [REACTOME]
A
B
Cell type
Disease model
ACKR2
ACKR3
ACKR4
CCL11
CCL16
CCL17
CCL19
CCL2
CCL20
CCL21
CCL22
CCL25
CCL28
CCL3
CCL5
CCL7
CCR1
CCR2
CCR3
CCR5
CCR6
CCR7
CCR9
CCRL2
CX3CL1
CX3CR1
CXCL1
CXCL10
CXCL11
CXCL12
CXCL13
CXCL16
CXCL2
CXCL3
CXCL5
CXCL6
CXCL8
CXCL9
CXCR1
CXCR2
CXCR3
CXCR4
CXCR5
CXCR6
PF4
PPBP
XCL1
XCL2
XCR1
Cell type
Dendritic Cells
Keratinocytes
Langerhans Cells
Melanocytes
T-cell
Disease model
IMQ
OVA
OXA
Orthology status
Not ortholog
FC significance
FDR <= 0.05

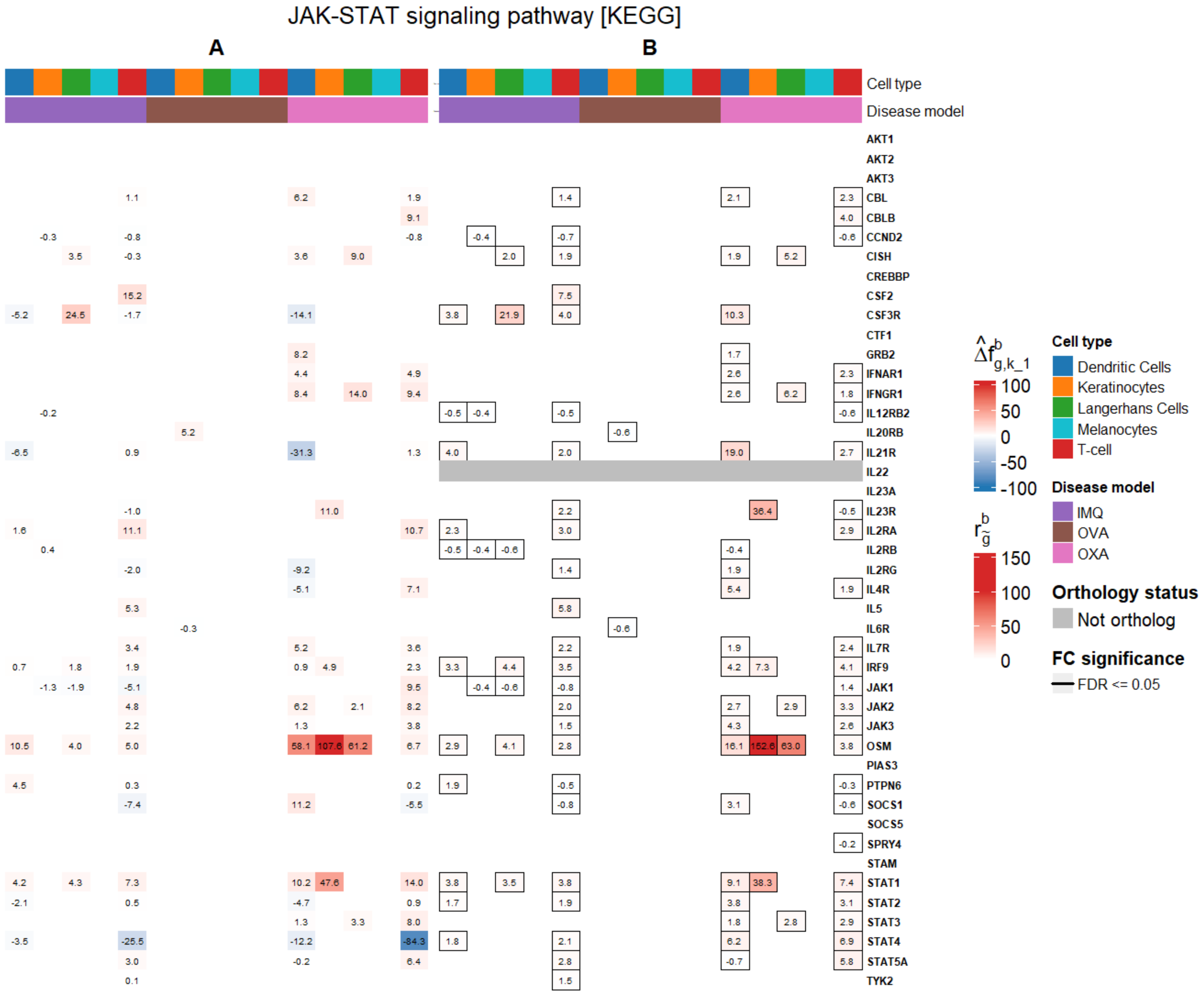
JAK-STAT signaling pathway [KEGG]
A
B
Cell type
Disease model
AKT1
AKT2
AKT3
CBL
CBLB
CCND2
CISH
CREBBP
CSF2
CSF3R
CTF1
GRB2
IFNAR1
IFNGR1
IL12RB2
IL20RB
IL21R
IL22
IL23A
IL23R
IL2RA
IL2RB
IL2RG
IL4R
IL5
IL6R
IL7R
IRF9
JAK1
JAK2
JAK3
OSM
PIAS3
PTPN6
SOCS1
SOCS5
SPRY4
STAM
STAT1
STAT2
STAT3
STAT4
STAT5A
TYK2
$\hat{\Delta f}^{b}_{g,k\_1}$
100
50
0
-50
-100
$r^{b}_{\tilde{g}}$
150
100
50
0
Cell type
Dendritic Cells
Keratinocytes
Langerhans Cells
Melanocytes
T-cell
Disease model
IMQ
OVA
OXA
Orthology status
Not ortholog
FC significance
FDR <= 0.05

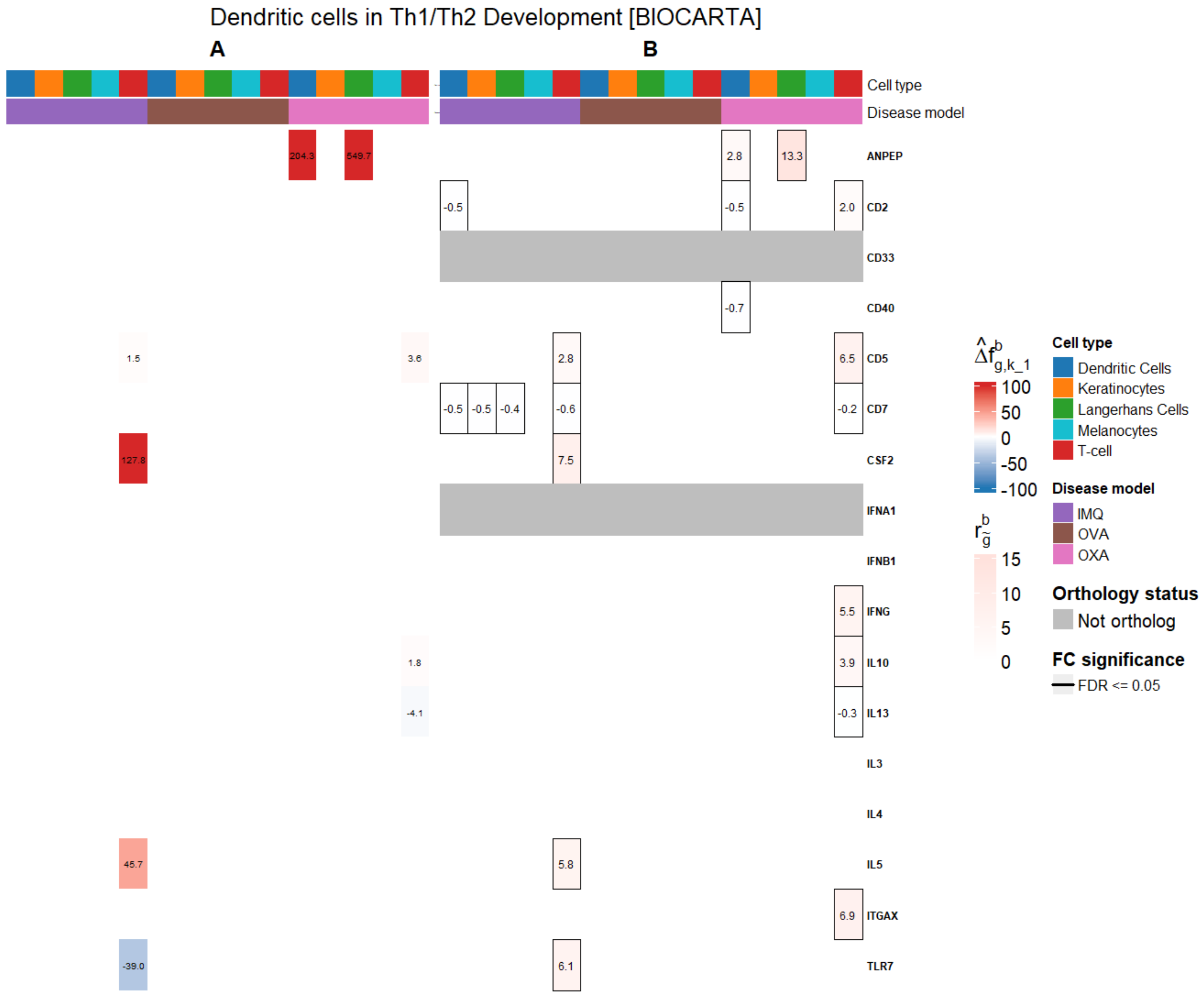


**Fig. S3. Gene contribution and disease model estimated $r^b_{\tilde{g}}$. A)** Gene contribution to cell type recapitulation by disease model. If gene set size of pathway is greater than 50, only the top 20 contributing genes, for each cell type, were displayed. Blank gene contributions correspond to 0 values. **B)** Computed $r^b_{\tilde{g}}$ by disease model. Grey FC refer to genes without one-to-one ortholog and/or not sequenced in disease model. Framed FC refer to statistically significant $FDR \leq 0.05$ genes, as per FindMarkers. Blank FC correspond to 0 values.

**Table S2.** References of Manuscript's Table 2

| Pathway | Cell type | Top 5 genes | Reference direction |
|---|---|---|---|
| JAK-STAT signaling pathway [KEGG] | T-cell | IL13, IL26, IL2RA, IL7, IFNGR1 | (Napolitano et al., 2023), (Kamijo et al., 2020), (Jia et al., 2023) (Park et al., 2023), (Wasserer et al., 2024) |
| | Dendritic Cell | IL23A, SPRED1,SOCS1, IFNL1, OSM | (Napolitano et al., 2021), (Sakai et al., 2022), (Kopalli et al., 2022) (Philip et al., 2024), (Suehiro et al., 2023) |
| | Langerhan Cell | IL22RA2, CCND2, CCND1, JAK1, STAT6 | (Bangert et al., 2024), NA, NA, (Huang et al., 2022), (Antczak et al., 2016) |
| | Keratinocyte | CCND3, SPRY1, IL15RA, IFNAR2, IL15 | NA, (Cui et al., 2024), (Jones et al., 2016) (Morizane et al., 2023), (Karlen et al., 2020) |
| | Melanocyte | CCND3, IFNGR2, CCND2, IL10RA, CCND1 | (Alekseenko et al., 2010), (Su et al., 2020), NA (Zhou et al., 2016), (Alekseenko et al., 2010) |
| Dendritic Cells in Th1/Th2 development [BIOCARTA] | T-cell | IL13, IL5, CSF2, TLR7 | (Napolitano et al., 2023), (Antosz et al., 2024), [(Mu et al., 2021), (Xing et al., 1997)], (Jeisy-Scott et al., 2011) |
| | Dendritic Cell | ANPEP, CSF2, IL13 | (Lu et al., 2020), (Taha et al., 1998), (Lamiable et al., 2022) |
| | Langerhan Cell | ANPEP | (Lu et al., 2020) |
| | Keratinocyte | ITGAX | (Zhong et al., 2021) |
| | Melanocyte | IL10, ITGAX, CD7, CD33 | (Ohmen et al., 1995), (Zhong et al., 2021), NA, NA |
| Cytokine-Cytokine receptor interaction [KEGG] | T-cell | IL13, CCR2, TNFSF10, CXCL13, IL1R2 | (Napolitano et al., 2023), [(Bakos et al., 2017), (Nedoszytko et al., 2014)], (Vassina et al., 2005), (Kwon et al., 2021), (Yamamoto-Hanada et al., 2023) |
| | Dendritic Cell | IL23A, CCR6, CCL5, CCL3L1, TNFRSF14 | (Napolitano et al., 2021), (Gros et al., 2009), (Tsoi et al., 2020) (Alkon et al., 2022), (Bangert et al., 2021) |
| | Langerhan Cell | IL22RA2, PLEKHO2, IL23A, IL7R, CCR1 | (Bangert et al., 2024), (Nousbeck et al., 2022), (Napolitano et al., 2021) (Gonzalez-Rodriguez et al., 2022), (Nedoszytko et al., 2014) |
| | Keratinocyte | IL15RA, TNFRSF12A, CCR2, TNFRSF11A, IFNAR2 | (Morizane et al., 2023), (Bangert et al., 2021), (Nedoszytko et al., 2014) NA, (Hile et al., 2020) |
| | Melanocyte | CX3CL1, TGFB2, IFNGR2, TNFSF13B, IL10RA | (Staumont-Sallé et al., 2014), (Arkwright et al., 2001), (Su et al., 2020) NA, (Zhou et al., 2016) |
| Chemokine receptors bind chemokines [REACTOME] | T-cell | CCR2, CXCL13, CCR1, CXCL8, CCR7 | [(Bakos et al., 2017), (Nedoszytko et al., 2014)], (Kwon et al., 2021), (Nedoszytko et al., 2014) (Morgner et al., 2023), (Bangert et al., 2021) |
| | Dendritic Cell | CCL5, CCR6, CCL3L1, CCL13, CXCL2 | (Tsoi et al., 2020), (Gros et al., 2009), (Alkon et al., 2022) (Bangert et al., 2021), NA |
| | Langerhan Cell | CCR1, CCL17, CXCR4, CCR10, CCRL2 | (Nedoszytko et al., 2014), (Bangert et al., 2021), (Dubrac et al., 2010) NA, (Nousbeck et al., 2022) |
| | Keratinocyte | CCR2, CXCL2, CCL7, XCR1, CCL20 | (Nedoszytko et al., 2014), NA, (Gros et al., 2009), NA, (Nakayama et al., 2001) |
| | Melanocyte | CX3CL1, CXCL8, CXCR6 | (Staumont-Sallé et al., 2014), , (Zhang et al., 2023) |
