## Supplementary Material S4 for "singIST: an integrative method for comparative single-cell transcriptomics between disease models and humans"

| Subject ID | Age | Sex | Ethnicity | EASI | Diagnosis | Sample type | scRNA-seq protocol | GEO identifier | Reference |
|---|---|---|---|---|---|---|---|---|---|
| AD1 | 18 | male | Caucasian | 34.2 | AD | Suction blister | 10X Genomics | GSM4653855 | Rojahn et.al 2020 |
| AD2 | 19 | female | Caucasian | 44.6 | AD | Suction blister | 10X Genomics | GSM4653856 | Rojahn et.al 2020 |
| AD3 | 34 | male | Caucasian | 44.7 | AD | Suction blister | 10X Genomics | GSM4653857 | Rojahn et.al 2020 |
| AD4 | 33 | female | Caucasian | 5.5 | AD | Suction blister | 10X Genomics | GSM4653858 | Rojahn et.al 2020 |
| AD13 | 45 | female | Caucasian | 27.4 | AD | Suction blister | 10X Genomics | GSM4800173 | Bangert et.al 2021 |
| HC1-2 | 42 | female | Caucasian | n.a. | | Suction blister | 10X Genomics | GSM4653863<br>GSM4653864 | Rojahn et.al 2020 |
| HC3 | 47 | female | Caucasian | n.a. | | Suction blister | 10X Genomics | GSM4653865 | Rojahn et.al 2020 |
| HC4 | 49 | female | Caucasian | n.a. | | Suction blister | 10X Genomics | GSM4653866 | Rojahn et.al 2020 |
| HC5 | 39 | male | Caucasian | n.a. | | Suction blister | 10X Genomics | GSM4653867 | Rojahn et.al 2020 |

| Samples | Group | Group description | Organism | Strain | Sex | Age | Tissue | Sample type | scRNA-seq protocol | GEO identifier | Reference |
|---|---|---|---|---|---|---|---|---|---|---|---|
| IMQ-C1 | IMQ-C | no IMQ treatment negative control (Vanicream) | Mus musculus | C57BL/6J | Male | 9 weeks | ear | skin biopsy | 10X Genomics | GSM4490718 | Liu et.al 2020 |
| IMQ-C2 | IMQ-C | no IMQ treatment negative control (Vanicream) | Mus musculus | C57BL/6J | Male | 9 weeks | ear | skin biopsy | 10X Genomics | GSM4490719 | Liu et.al 2020 |
| IMQ-C3 | IMQ-C | no IMQ treatment negative control (Vanicream) | Mus musculus | C57BL/6J | Male | 9 weeks | ear | skin biopsy | 10X Genomics | GSM4490720 | Liu et.al 2020 |
| IMQ1 | IMQ | Imiquimod 5% cream | Mus musculus | C57BL/6J | Male | 9 weeks | ear | skin biopsy | 10X Genomics | GSM4490721 | Liu et.al 2020 |
| IMQ2 | IMQ | Imiquimod 5% cream | Mus musculus | C57BL/6J | Male | 9 weeks | ear | skin biopsy | 10X Genomics | GSM4490722 | Liu et.al 2020 |
| IMQ3 | IMQ | Imiquimod 5% cream | Mus musculus | C57BL/6J | Male | 9 weeks | ear | skin biopsy | 10X Genomics | GSM4490723 | Liu et.al 2020 |
| OXA-C1 | ETOH | control ethyl alcohol (EtOH) | Mus musculus | C57BL/6J | Male | 10 weeks | ear | skin biopsy | 10X Genomics | GSM4490724 | Liu et.al 2020 |
| OXA-C2 | ETOH | control ethyl alcohol (EtOH) | Mus musculus | C57BL/6J | Male | 10 weeks | ear | skin biopsy | 10X Genomics | GSM4490725 | Liu et.al 2020 |
| OXA-C3 | ETOH | control ethyl alcohol (EtOH) | Mus musculus | C57BL/6J | Male | 10 weeks | ear | skin biopsy | 10X Genomics | GSM4490726 | Liu et.al 2020 |
| OXA1 | OXA | Oxazolone challenged | Mus musculus | C57BL/6J | Male | 10 weeks | ear | skin biopsy | 10X Genomics | GSM4490727 | Liu et.al 2020 |
| OXA3 | OXA | Oxazolone challenged | Mus musculus | C57BL/6J | Male | 10 weeks | ear | skin biopsy | 10X Genomics | GSM4490728 | Liu et.al 2020 |
| OXA3 | OXA | Oxazolone challenged | Mus musculus | C57BL/6J | Male | 10 weeks | ear | skin biopsy | 10X Genomics | GSM4490729 | Liu et.al 2020 |
| EC-OVA-1 | OVA | ovalbumine epicutaneously sensitized | Mus musculus | Balb/c | n.a | n.a | ear | skin biopsy | 10X Genomics | GSM5831751 | Leyva-Castillo et.al 2022 |
| EC-OVA-2 | OVA | ovalbumine epicutaneously sensitized | Mus musculus | Balb/c | n.a | n.a | ear | skin biopsy | 10X Genomics | GSM5831752 | Leyva-Castillo et.al 2022 |
| EC-OVA-3 | OVA | ovalbumine epicutaneously sensitized | Mus musculus | Balb/c | n.a | n.a | ear | skin biopsy | 10X Genomics | GSM5831753 | Leyva-Castillo et.al 2022 |
| EC-SAL-1 | SAL | saline solution epicutenously sensitized | Mus musculus | Balb/c | n.a | n.a | ear | skin biopsy | 10X Genomics | GSM5831748 | Leyva-Castillo et.al 2022 |
| EC-SAL-2 | SAL | saline solution epicutenously sensitized | Mus musculus | Balb/c | n.a | n.a | ear | skin biopsy | 10X Genomics | GSM5831749 | Leyva-Castillo et.al 2022 |
| EC-SAL-3 | SAL | saline solution epicutenously sensitized | Mus musculus | Balb/c | n.a | n.a | ear | skin biopsy | 10X Genomics | GSM5831750 | Leyva-Castillo et.al 2022 |
